# The creation and validation of a fully animal component-free media for select adherent cell types

**DOI:** 10.1101/2025.02.05.636679

**Authors:** Nicolette B. Mogilever, Marie-Hélène Godin Pagé, Anjolaoluwatikiitan Solola, Andrew E. Pelling

**Affiliations:** Department of Biology, Gendron Hall, 30 Marie Curie, University of Ottawa, Ottawa, ON, K1N5N5 Canada; Division of Experimental Medicine, Department of Medicine, 1001 Decarie Blvd, McGill University, Montreal, QC, H4A3J1, Canada; Department of Biomedical Engineering, Weill Hall, 237 Tower Rd, Cornell University, Ithaca, NY, 14850, USA; Department of Physics, STEM Complex, 150 Louis Pasteur Pvt., University of Ottawa, Ottawa, ON, K1N5N5 Canada

**Author notes:** Author for correspondence: Andrew E. Pelling, University of Ottawa.

## Abstract

Fetal Bovine Serum (FBS) is one of the most commonly used media supplement for the maintenance of mammalian cell types, yet the expensive costs, ethical concerns, and lot-to-lot variation have provoked a clear need for a serum that is standardized and derived from non-animal sources. Several serum-free formulations have been developed in the past, however they are often cell type specific, contain animal-derived components, and lack long-term culture validation. In this study, we developed a novel animal component-free (ACF) media and investigated its effectiveness on four commonly used mammalian cell lines via long-term (up to 90 days) morphological, transcriptomic, and proliferative analyses. Cells cultured in our ACF medium exhibited comparable cellular morphologies and equal or greater growth rates compared with cells cultured with FBS. Additionally, differentially expressed genes between the FBS-grown and ACF-grown groups were predominantly associated with functions linked to proliferation and cell attachment. The findings from this study indicate that this medium is a suitable replacement to FBS-containing medium for several common cell lines.

## Introduction

Cell culture systems, pivotal for studying physiological, biological, and pharmacological activities at the cellular level, have significantly advanced biology with broad applications ranging from drug evaluation and tissue engineering to vaccine production and cultivated meats [1–4]. Cell culture medium, the vehicle that supports cell viability, is essential to any cell culture in vitro [5,6]. The proteins, vitamins, minerals, hormones, growth factors, and other key nutrients that are responsible for providing this support to mammalian cells must be supplied by the medium they are grown in [7,8]. The major supplier for these essential components has been commonly derived from animal sera, with fetal bovine serum (FBS) being the alleged ‘gold standard’ supplement since its introduction to cell culture scientists in 1958 [9,10]. Other mammalian-derived sera from chicken, sheep, pig, turkey, and horse blood can also be used in certain protocols or for specific cell types, but FBS continues to be the most widely used supplement [11]. A by-product of the animal agriculture industry, its production varies based on changes in the public’s beef consumption, outbreaks of bovine-related diseases, and changes in crop prices for herd feed [12]. The high demand for meat products has led to large cattle inventories and therefore it is unsurprising that FBS can be relatively easily sourced and distributed to scientists worldwide [12,13]. Although practical, FBS carries several ethical, safety, and reproducibility concerns and ideally a more sophisticated, controllable and reproducible alternative should be developed [14,15].

The high degree of lot-to-lot variability in FBS can be one significant cause of irreproducibility in research findings from lab to lab around the world [14–17]. This has a detrimental impact on academic science and on the commercialization of biotechnologies aimed at producing consumer products, cell-based therapeutics, and medical assays, and it is not fully understood how each component contributes to either proliferation or differentiation, at what capacities, and in differing cell types [18–20]. Chemically-defined, serum-free media (SFM) alternatives pose fewer ethical concerns, eliminate the potential of biological contaminants, create more controlled culture conditions, have more stable costs and could be less expensive altogether [14,21,22]. Many literature reviews on SFM development methodologies and essential components have been conducted [14,15,18–20]. Commercially available and open-source serum supplements have been produced previously with over 100 different serum-free formulations to date, however they are typically optimized for the growth of specific cell types [13–15], for example chondrocytes [25–29], myoblasts [30–33], satellite cells [34–37], HeLa cells [38–40], Vero cells [41–45], and a variety of fibroblastic lines [46–51]. It is important to note that although there is some overlap in the components used by these groups, the implemented concentrations and ingredients vary between most of the developed formulations and the quantitative assessment methods used to validate each formulation differ greatly from one another, making comparisons difficult.

These alternative media additives provide a strong starting point for the development of a more generalized serum replacement. The goal of the research presented here is to develop and validate a general-use, well-characterized, ACF serum supplement to address the obstacles described above, for several different adherent cell lines. While SFM eliminate the use of whole serum, they may still contain animal-derived components such as bovine serum albumin or animal-derived growth factors; in contrast, ACF media are explicitly formulated without any animal-derived ingredients, offering greater consistency, ethical advantages, and regulatory alignment for clinical and industrial applications [14,18,23]. The ACF media we developed was validated by looking at proliferation and morphological analyses as well as transcriptomics in long-term culture. Herein we demonstrate the creation of a synthetic and recombinant component-based media that is validated for the NIH 3T3 fibroblast, C2C12 myoblast, MC3T3 pre-osteoblast, and HFF human foreskin fibroblast cell lines, which can be utilized, and independently validated by scientists worldwide who depend on mammalian cell culture processes. The experiments carried out here provide the cell culture community with verification of a functional serum replacement and create a modern starting point for non-computational serum-free fabrication.

## Results

### Serum-free media creation

A multi-step methodology was carried out in order to investigate various combinations of supplements for cell culture media. We examined their interactions and fine-tuned their concentrations to create a SFM formulation suitable for supporting the growth of NIH 3T3 fibroblasts, and later made minor adjustments of the formulation for 3 commonly used cell lines in research. To do so, the following steps were carried out:

1. By assessing the studies and reviews that investigated serum-free media formulations, essential components to adherent mammalian cell culture of widely used cell lines were identified [14–51]. From these reviews and primary research articles, a database was created to summarize any components that were used in successful serum-free culture as well as their concentrations. By adapting the recommendations from van der Valk’s 2010 review, the essential components needed to begin formulating SFM are presented (Fig. 1). We examined the components which overlapped studies most and then chose the ones that could be made ACF to start in our first formulation. Initially, this consisted of a strongly recommended basal medium (DMEM/F-12), 2 hormones (insulin, prostaglandin E2), 2 carrier proteins (transferrin, albumin), 1 lipid derivative (ethanolamine), and 3 growth factors (epidermal growth factor (EGF), fibroblast growth factor (FGF2), transforming growth factor beta (TGFb).
2. With this initial formulation, NIH 3T3s were sequentially adapted from serum-containing medium to serum-free medium over a 10-day period *(see Methods, Adaptation and Serum-free Medium).* Next, cell viability using trypan blue exclusion was measured after seven days of culture. Although this initial formulation allowed for high viability (>90%) after seven days, the morphology of the cells as shown using phase contrast microscopy exhibited much more elongated, spindle-like morphology as compared to the cells grown in FBS-containing medium. Thus, further optimization of the components was required.
3. The concentrations of four components were first chosen to be modified including the 3 growth factors (EGF, FGF2, TGFb) and 1 hormone (Prostaglandin E2, PGE2). The reason that these factors were chosen to be modified initially while keeping the others constant was because the other components showed much more consistency across serum-free media studies both in their implementation and concentration, while these growth factors and hormone were implemented quite differently across successful serum-free research. In the past, matrices for culture media components have been used to identify the most significant factors that influence a certain outcome [150,151]. In this study, we modified this approach by setting up a matrix that allowed us to screen for different combinations of concentrations of these four chosen media components. We implemented this using a non-statistical approach with 2 desired outcomes: a) maintaining high viability (>90%) at the seventh day of culture, and b) exhibiting a similar morphological pattern via phase contrast microscopy as compared to cells grown in FBS-containing media. Table 1 summarizes all concentrations and outcomes found in this optimization study.
4. From this matrix, we found several combinations that allowed for our two desired outcomes. Next, to create simplicity and minimize costs downstream, we chose the formulation that used the minimal concentration of growth factors but still fostered the same outcomes, since growth factors are often responsible for most of the high serum-free media costs.
5. Finally, this formulation was used to quantitatively assess long-term proliferation in NIH 3T3 fibroblasts and HFF human foreskin fibroblasts. This final formulation did not have optimal success with the other 2 cell lines used in this study (C2C12 myoblasts, MC3T3 osteoblasts) as their proliferation rates were lower and morphologies too dissimilar from their FBS-grown counter parts. As a result, two media components were altered for these cell lines. Firstly, the basal medium was changed to their native basal medium that is most commonly used for their culture – DMEM for C2C12 and aMEM for MC3T3 cells.

**Figure 1:**
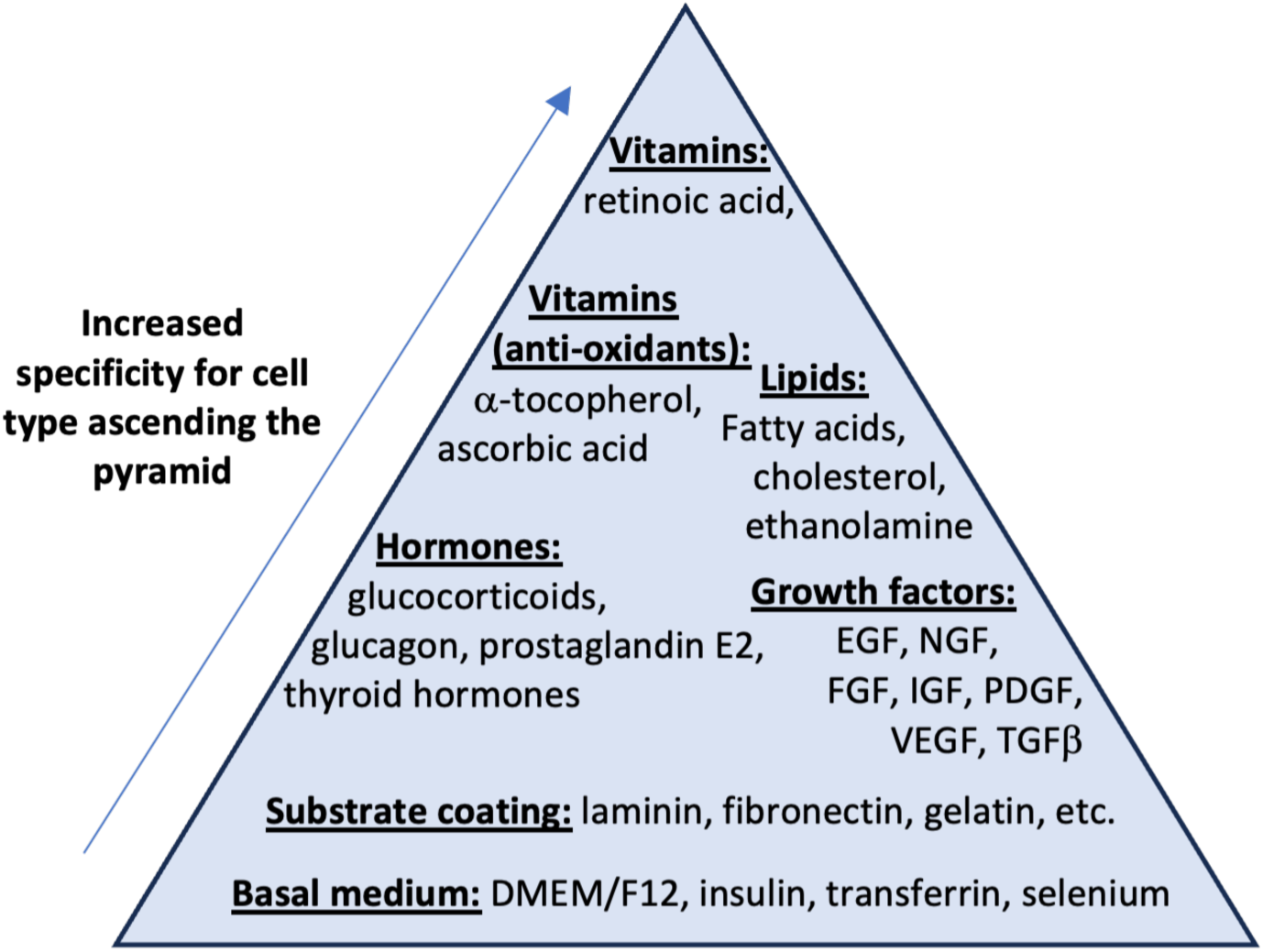
Schematic modular approach for the development of serum-free media. Start the formulation at the base of the pyramid and move upwards to increase specificity for the desired cell line. Each heading provides examples of the types of components that can be added but is not a comprehensive list. *Adapted from van der Valk, 2010.* [23].

**Table 1:**
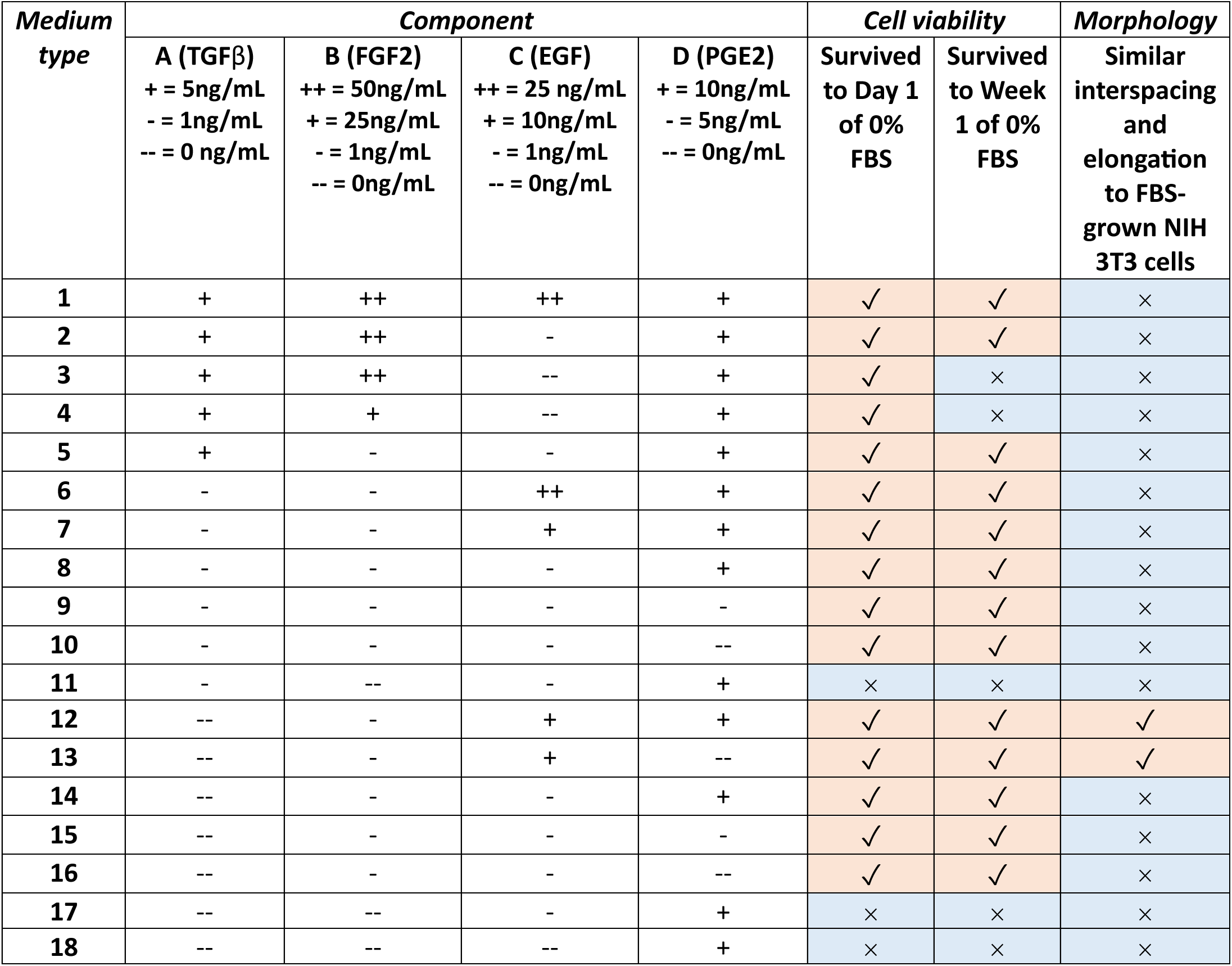
Optimization matrix for final ACF medium formulation. Components A, B, C, and D were chosen based off the literature suggesting that these growth factors and hormones may be related to more volatile changes in cell growth. Their concentrations designated as (++) represent the high end of the given range of concentrations based off the preliminary medium formulation. Their concentrations are designated as (-) represent the low end of the given range of concentrations based off pre-existing literature. Their concentrations designated as (--) represent a concentration of zero. Each of the 18 media formulations were allowed to proliferate for at least 7 days since any exposure to FBS, if possible.

| Medium type | Component |  |  |  | Cell viability |  | Morphology |
| --- | --- | --- | --- | --- | --- | --- | --- |
|  | A (TGFβ)<br>+ = 5ng/mL<br>- = 1ng/mL<br>-- = 0 ng/mL | B (FGF2)<br>++ = 50ng/mL<br>+ = 25ng/mL<br>- = 1ng/mL<br>-- = 0ng/mL | C (EGF)<br>++ = 25 ng/mL<br>+ = 10ng/mL<br>- = 1ng/mL<br>-- = 0ng/mL | D (PGE2)<br>+ = 10ng/mL<br>- = 5ng/mL<br>-- = 0ng/mL | Survived to Day 1 of 0% FBS | Survived to Week 1 of 0% FBS | Similar interspacing and elongation to FBS-grown NIH 3T3 cells |
| 1 | + | ++ | ++ | + | ✓ | ✓ | × |
| 2 | + | ++ | - | + | ✓ | ✓ | × |
| 3 | + | ++ | -- | + | ✓ | × | × |
| 4 | + | + | -- | + | ✓ | × | × |
| 5 | + | - | - | + | ✓ | ✓ | × |
| 6 | - | - | ++ | + | ✓ | ✓ | × |
| 7 | - | - | + | + | ✓ | ✓ | × |
| 8 | - | - | - | + | ✓ | ✓ | × |
| 9 | - | - | - | - | ✓ | ✓ | × |
| 10 | - | - | - | -- | ✓ | ✓ | × |
| 11 | - | -- | - | + | × | × | × |
| 12 | -- | - | + | + | ✓ | ✓ | ✓ |
| 13 | -- | - | + | -- | ✓ | ✓ | ✓ |
| 14 | -- | - | - | + | ✓ | ✓ | × |
| 15 | -- | - | - | - | ✓ | ✓ | × |
| 16 | -- | - | - | -- | ✓ | ✓ | × |
| 17 | -- | -- | - | + | × | × | × |
| 18 | -- | -- | -- | + | × | × | × |

And lastly, after similar optimization steps described above, the concentration of FGF2 increased from 1 ng/mL to 30 ng/mL for both cell lines, TGFb was kept as is, and PGE2 was eliminated altogether. Separate from the medium formulation, a substrate coating was added to increase cell adhesion in all cell lines – the details of this finding and execution are detailed in the Methods *(Animal- and plant-based substrate coatings*). The final medium formulation used for all cell types is summarized in Table 2. This final iteration of the ACF formulation is the medium used in the treatment groups in the following presented results.

**Table 2:**
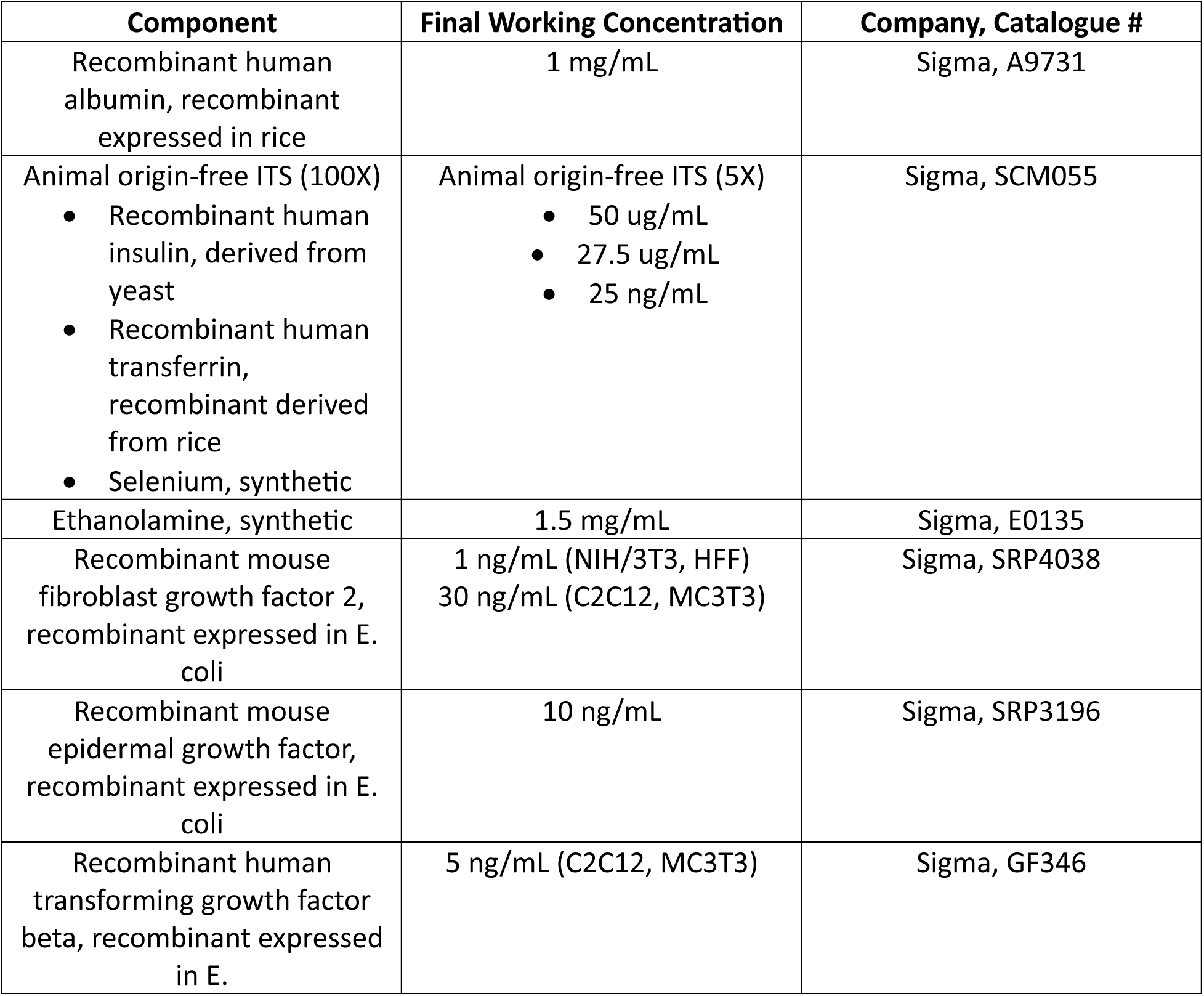
Final optimized formulation for animal component-free medium. Final recipe for optimized medium formulation used in this study. These components were added in supplement to the same basal medium as used in the FBS-grown control groups (*see Methods for basal medium description*) with 1% Penicillin and Streptomycin.

| **Component** | **Final Working Concentration** | **Company, Catalogue #** |
| --- | --- | --- |
| Recombinant human albumin, recombinant expressed in rice | 1 mg/mL | Sigma, A9731 |
| Animal origin-free ITS (100X)<br>• Recombinant human insulin, derived from yeast<br>• Recombinant human transferrin, recombinant derived from rice<br>• Selenium, synthetic | Animal origin-free ITS (5X)<br>• 50 ug/mL<br>• 27.5 ug/mL<br>• 25 ng/mL | Sigma, SCM055 |
| Ethanolamine, synthetic | 1.5 mg/mL | Sigma, E0135 |
| Recombinant mouse fibroblast growth factor 2, recombinant expressed in E. coli | 1 ng/mL (NIH/3T3, HFF)<br>30 ng/mL (C2C12, MC3T3) | Sigma, SRP4038 |
| Recombinant mouse epidermal growth factor, recombinant expressed in E. coli | 10 ng/mL | Sigma, SRP3196 |
| Recombinant human transforming growth factor beta, recombinant expressed in E. | 5 ng/mL (C2C12, MC3T3) | Sigma, GF346 |

### ACF media supports fibroblast proliferation and viability

After sequential media adjustment, NIH 3T3s were inoculated onto tissue culture plates and cultured for 7 days. At days 1, 4 and 7 cell concentration was quantified in an automated cell counter (Fig. 2A). On days 1 and 4 there was no difference in cell concentration (P=0.4468 and P=0.0776, respectively), however on the seventh day of culture, the ACF medium resulted in a significantly lower cell concentration compared to the FBS medium group (P=0.0362).

**Figure 2:**
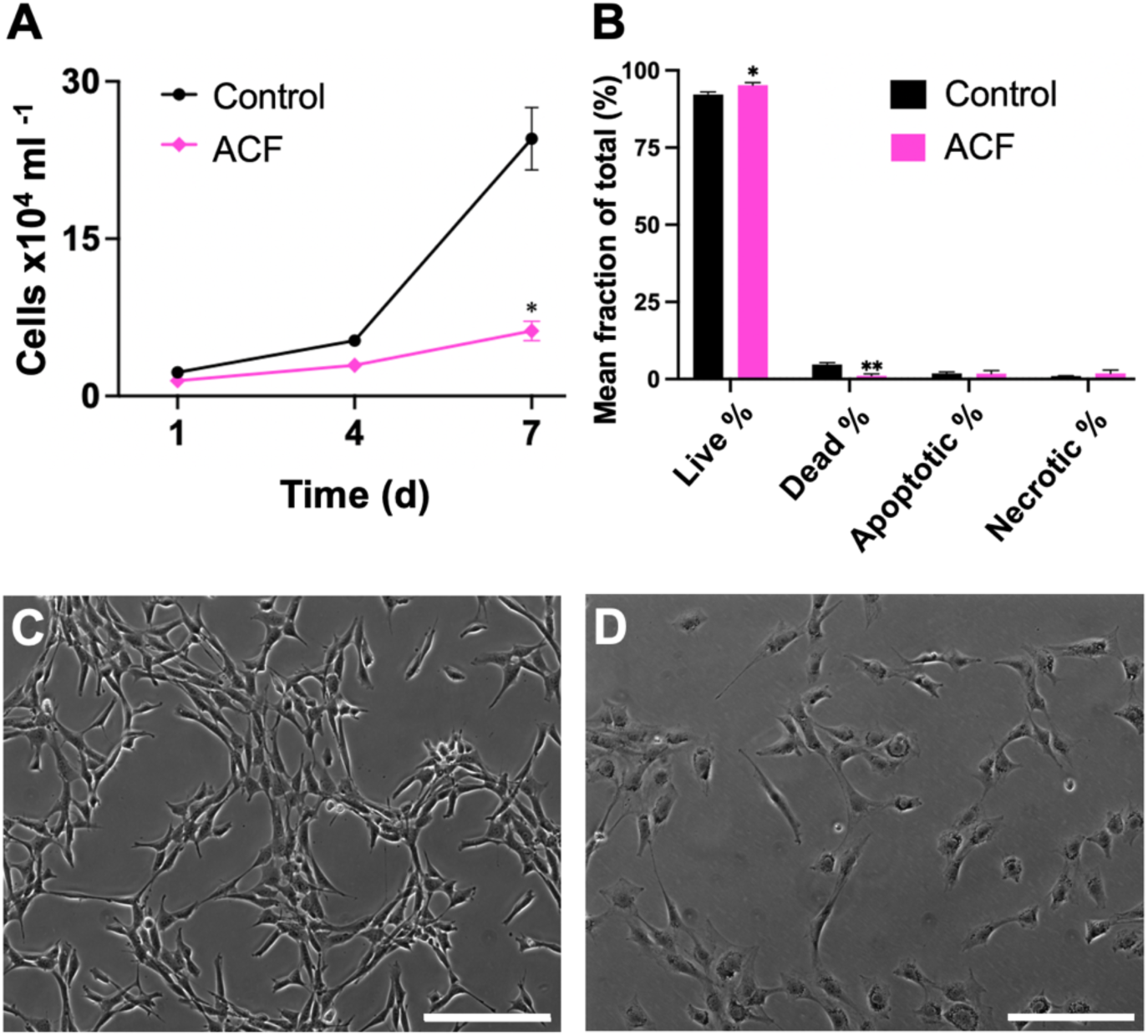
Short-term (<7 day) cell viability and morphology of NIH 3T3 fibroblasts. **(A)** Mean concentration of cells (x10^4^ per ml) after 1, 4, and 7 days since initial seeding. **(B)** Mean live, dead, early apoptotic, and necrotic cell populations represented as a percent of the total cell concentration, measured by Annexin-V/PI staining. N=3 for each time point and medium condition. Each biological replicate is the mean of 3 technical replicates. Data are presented as mean ± SEM where *P<0.033, **P<0.002, ***P<0.001, as compared to the control medium. Most representative phase contrast images. Scale bars, 200um. **(C)** Control medium (10% FBS). **(D)** ACF medium (no coating).

Furthermore, Annexin-V/PI staining revealed that both ACF and FBS groups were able to maintain high cell viability (>80%) at each time point (Fig. 2B). Phase contrast images reveal significantly lower cell concentration in ACF vs FBS media on the seventh day of culture (Fig. 2C-D). Although qualitative, it was observed that cells in ACF medium tended to easily shear off the culture plates indicating that in the absence of FBS, cells were poorly adhered to the culture plate. Therefore, we conducted a series of experiments which explored the use of animal and plant-based surface coatings to solve this problem *(detailed in the Methods)*.

We included gelatin in our analyses as it is a widely used animal-derived coating that is known to increase adhesion in cell cultures [52–56]. We also chose a plant-based surface coating to use in our analyses as the goal of this work was to maintain the fully ACF quality of the culture system. Among the array of plant-based coatings applicable to adhesion in cell culture, soy stands out as a particularly promising candidate due to its unique combination of adhesive proteins, polysaccharides, and bioactive compounds [57–60]. Specifically, soy-derived proteins, such as soy protein isolate (SPI), have been recognized for their ability to form thin films on surfaces and promote cell adhesion [61–63]. These proteins possess adhesive domains that can interact with cell surface receptors, facilitating attachment to the substrate [64,65].

Furthermore, soy isoflavones, such as genistein and daidzein, have been reported to modulate cellular signaling pathways involved in adhesion and cytoskeletal organization [66–68]. These compounds can activate integrin-mediated signaling cascades, leading to enhanced focal adhesion formation and strengthening of cell-substrate interactions [68,69]. Thus we hypothesized that the application of a soy-derived coating to cell culture dishes could provide an adhesive microenvironment conducive to fibroblast attachment and spreading, similar to the effect that gelatin is widely used for. We developed a methodology to create a surface coating for Petri dishes from soy protein which was then used to support the adhesion of fibroblasts in ACF medium.

Cells were then cultured either on gelatin-coated dishes (ACF+gelatin) or on soy-coated dishes (ACF+soy). Cells grown in ACF+soy and ACF+gelatin showed similar total cell concentrations to the FBS-based group at days 1, 4, and 7. On the seventh day of culture, total cell concentrations in coated groups (ACF+gelatin and ACF+soy) were insignificantly changed from the FBS group (P=0.0850, P=0.0504, respectively) (Fig. 3A). All ACF media groups were able to proliferate (viability > 94%) by the seventh day of culture as determined by Annexin-V/PI staining (Fig. 3B). In order to view the morphology of the cells at these time points, phase contrast images were obtained. Fibroblasts grown in ACF+gelatin (Fig. 3E) and ACF+soy (Fig. 3F) showed similar densities (>75%) to the FBS group (Fig. 3C), consistent with the proliferation data. ACF+soy grown cells tended to grow in distinct clusters, similar to the morphology exhibited by the FBS group. Such recognizable clustering was not apparent in ACF+gelatin and the non-coated ACF groups.

**Figure 3:**
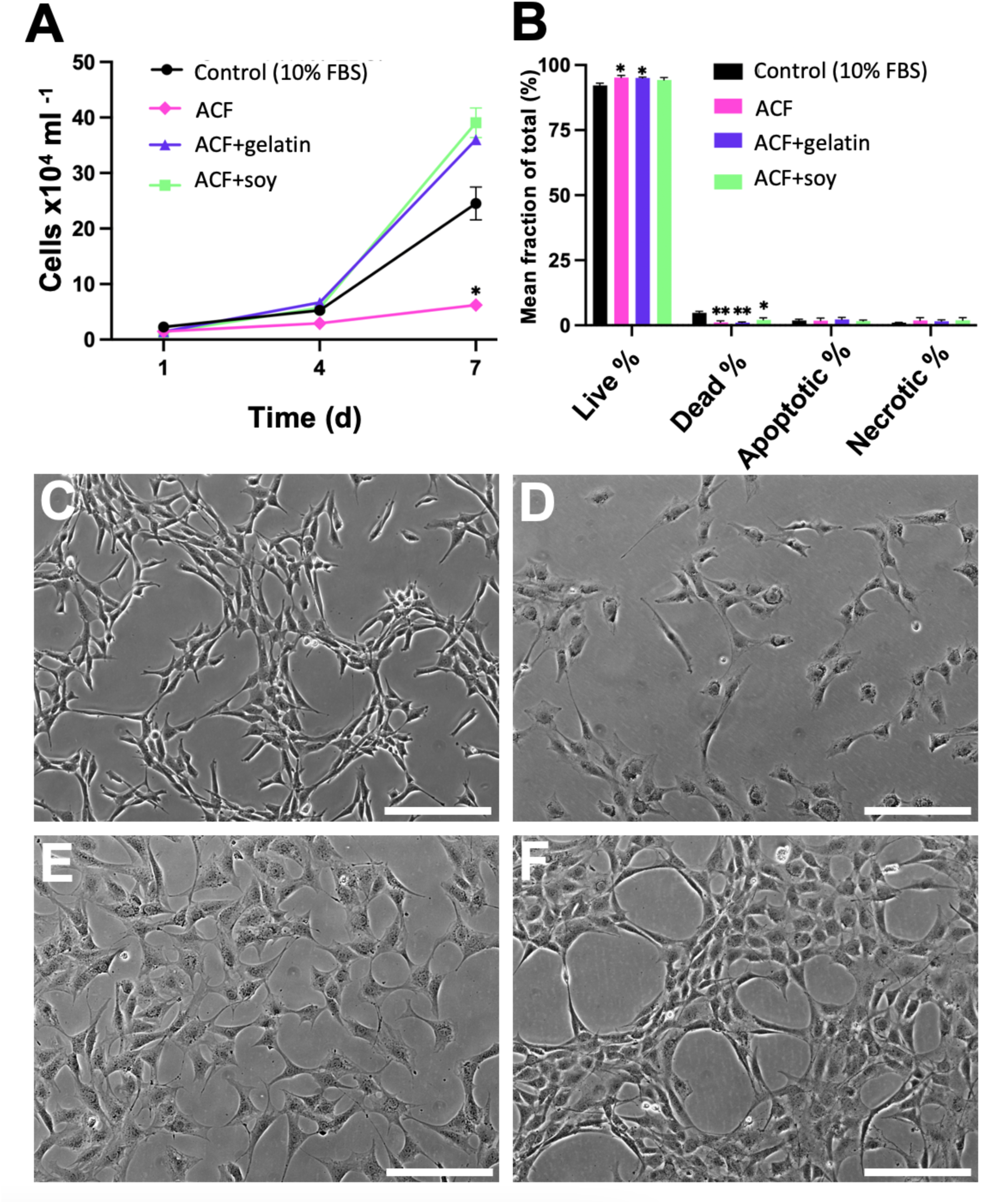
Short-term (<7 day) cell viability and morphology of NIH 3T3 fibroblasts in all ACF media conditions. **(A)** Mean concentration of cells (x10^4^ per ml). **(B)** Mean percentage of live, dead, early apoptotic, and necrotic cells as measured by Annexin-V/PI staining. N=3 for each time point and media condition. Each biological replicate is the mean of 3 technical replicates. Data are presented as mean ± SEM where *P<0.033, **P<0.002, ***P<0.001, as compared to the control medium. **(C-F)** Most representative phase contrast images. Scale bars, 200um. **(C)** Control medium (10% FBS). **(D)** ACF medium (no coating). **(E)** ACF medium (gelatin-coated). **(F)** ACF medium (soy-coated).

To better understand the long-term proliferative capabilities of the ACF medium on fibroblasts, we continued to culture the cells for up to 90 days. Every 30 days, assays for cell concentration and viability were performed, along with phase contrast microscopy. After 90 days in culture, all ACF conditions showed similar total cell concentrations to the FBS group at days 30 and 60 since full medium adjustment (Fig. 4A). On the 90^th^ day of culture, ACF+soy grown cells showed a significantly increased total cell concentration as compared to the FBS group (P=0.0040), while the ACF+gelatin and non-coated ACF groups showed insignificant changes in total cell concentration (P=0.0713, P=0.9854, respectively) (Fig. 4A). As shown by Annexin-V/PI staining, all ACF media groups were still viable after 90 days of culture (>95%) (Fig. 4B). Fibroblasts grown in ACF+gelatin medium showed a significantly increased live cell percentage as compared to the FBS group (P=0.004), while non-coated ACF and ACF+soy groups demonstrated an unchanged live cell percentage as compared to the FBS group. Phase contrast imaging demonstrates similar morphologies of ACF+soy cells to serum-grown cells, with non-coated and gelatin-coated ACF groups more elongated and less spread out (Fig. 4C-F).

**Figure 4:**
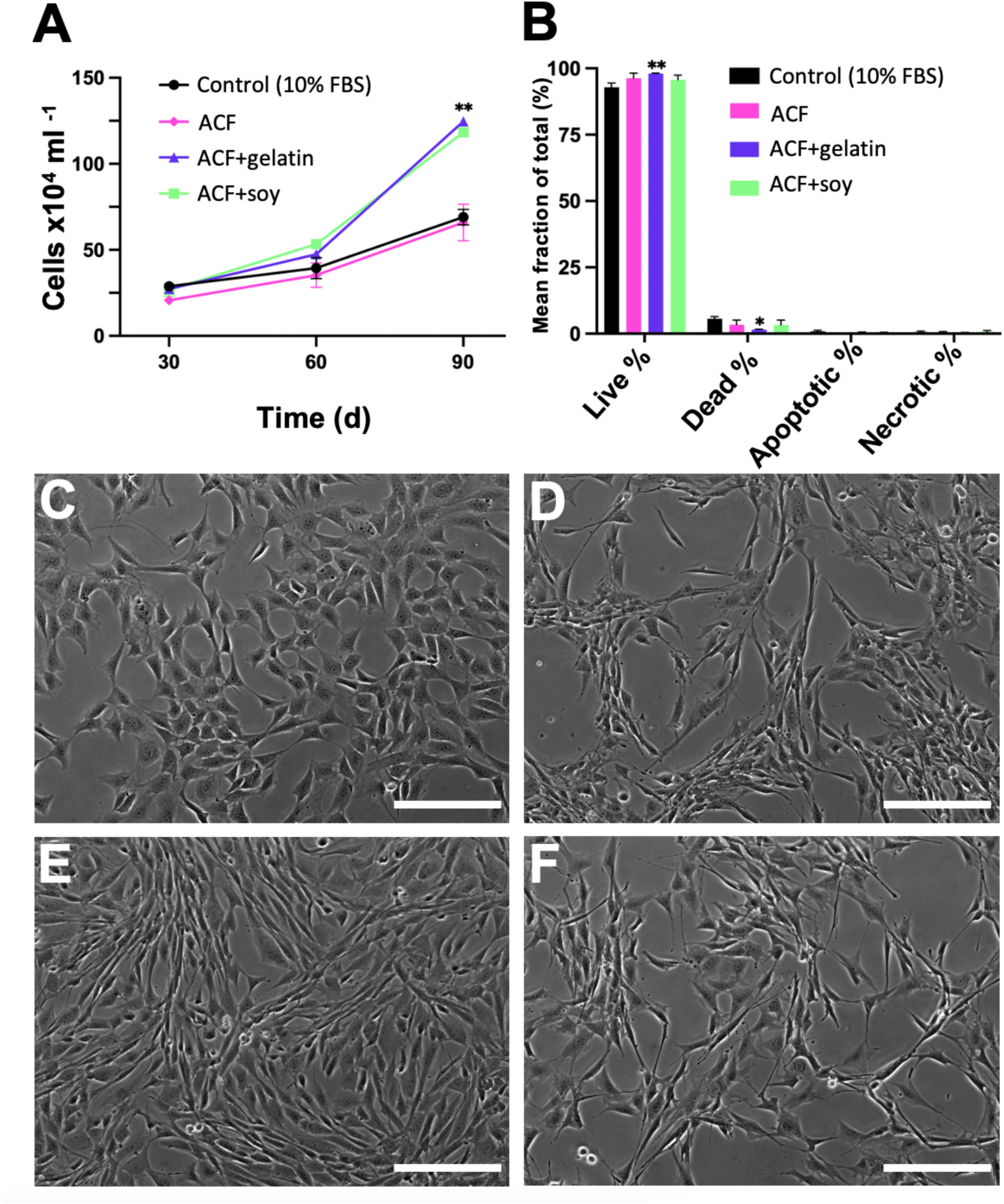
Long-term (30-90 day) cell viability and morphology of NIH 3T3 fibroblasts. (**A)** Mean concentration of cells (x10^4^ per ml). **(B)** Mean percentage of live, dead, early apoptotic, and necrotic cells as measured by Annexin-V/PI staining. N=3 for each time point and media condition. Each biological replicate is the mean of 3 technical replicates. Data are presented as mean ± SEM where *P<0.033, **P<0.002, ***P<0.001, as compared to the control medium. **(C-F)** Most representative phase contrast images. Scale bars, 200um. **(C)** Control medium (10% FBS). **(D)** ACF medium (no coating). **(E)** ACF medium (gelatin-coated). **(F)** ACF medium (soy-coated).

### Morphological differences of fibroblasts are minimal with ACF media

A major goal of this study was to not only create a serum-free alternative to FBS that can foster viable cells for short- and long-term periods but also that can allow typical cell lines used in research to behave similarly to those grown with FBS-based medium. When transitioning adherent cells from serum-containing to SFM, notable changes in cellular morphology are often observed [70–73]. Adherent cells cultured in SFM may exhibit alterations in their shape, size, and overall appearance compared to those cultured in serum-supplemented conditions [70–74]. These changes can be attributed to the absence of growth factors, hormones, and other serum components that play crucial roles in cell signaling and regulation of cellular processes [75–77]. Typically, cells cultured in SFM may display a more flattened or elongated morphology, with reduced cell-to-cell contact and altered cytoskeletal organization [78–80]. This morphological shift is often accompanied by changes in cellular behavior, such as decreased proliferation rates, alterations in gene expression profiles, and modifications in cell adhesion properties [81–84]. As a result, a morphological analysis on NIH 3T3s was performed to better understand how the ACF medium impacts cell size and shape as compared to cells grown in FBS-supplemented medium. The selection of nuclear and cell body areas as quantitative metrics that reflect morphology was driven by their relevance to fundamental cellular processes, including proliferation, differentiation, and migration, which are known to be influenced by changes in nutrient availability and signaling cues present in the culture medium [85–88]. This analytical strategy enabled us to discern subtle but significant differences in cellular morphology across distinct treatment groups, providing valuable insights into the cellular adaptations elicited by serum-free culture conditions.

Morphological analysis was performed via confocal laser scanning microscopy (CLSM) and fluorescent staining to analyze nuclear and cell body morphology for each culture condition. Cells in all groups demonstrated a typical fibroblastic morphology, with similar levels of interspaced, flattened cells (Fig. 5A-D). Image quantification allowed us to measure nuclear and cell body areas as well as their ratios (Fig. 5E-G). Similar mean nuclear areas of NIH 3T3s were found in the FBS-based medium (118.4±1.3μm) and in ACF+soy medium (114.5±1.1μm). These nuclear areas were found to be on average 24% smaller in uncoated ACF medium (90.2±1.0μm) and on average 30% larger in ACF+gelatin medium (152.7±1.6μm). Similar mean cell body areas of the cells were found in the FBS group (876.7±15.6μm) and in ACF+soy medium (854.1±17.5μm). The cell body areas were found to be on average 54% smaller in uncoated ACF medium (400.8±7.3μm) and on average 29% larger in ACF+gelatin medium (1129.7±20.4μm). Correspondingly, nuclear to cell area (N:C) ratios were calculated. Uncoated ACF medium demonstrated significantly different N:C ratios than the FBS group (P=0.0001), where no significant differences in N:C ratios obtained from the ACF+gelatin (P=0.9999) and ACF soy (P=0.9985) groups were found.

**Figure 5:**
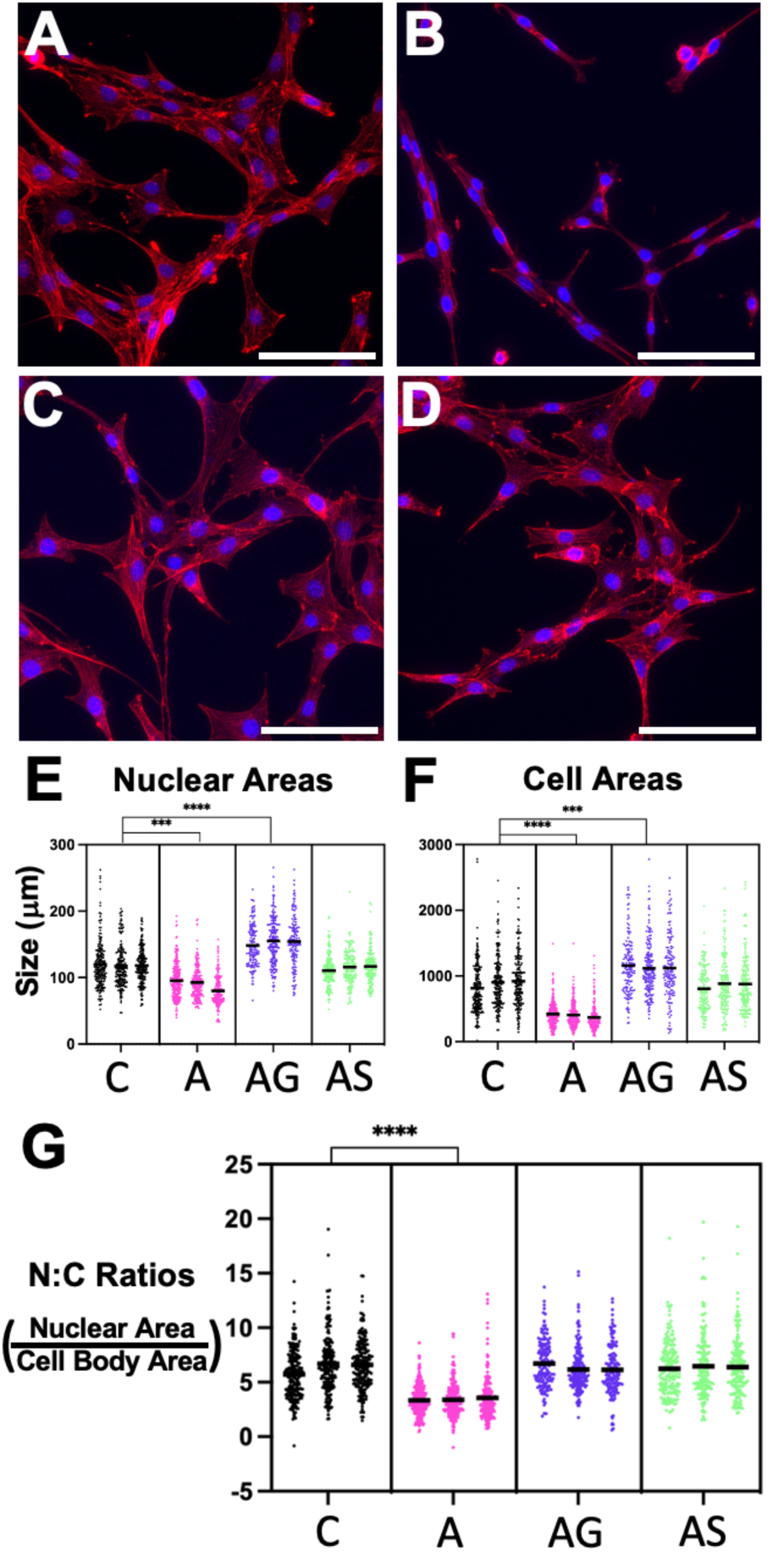
Morphological quantification of NIH 3T3 fibroblasts via cell sizing. **(A-D)** Representative confocal laser scanning microscopy images quantified after 7 days since initial seeding. Nuclei stained with DAPI (blue) and F-actin with Alexa Fluor^TM^ 647-Phalloidin (red). Scale bars, 100um. **(A)** Control medium (10% FBS). **(B**) ACF medium (no coating). **(C)** ACF medium (gelatin-coated). **(D)** ACF medium (soy-coated). **(E-G)** Beeswarm plots with nucleus & cell body areas as well as nucleus to cell body (N:C) ratios across media types (C=control, A=ACF no coating, AG=ACF+gelatin, AS=ACF+soy). Each point represents 1 cell. N=3 for each medium condition with 150+ technical replicates per biological replicate. Centered horizontal black lines represent the mean for each replicate. *P<0.033, **P<0.002, ***P<0.0002, ****P<0.0001 as compared to the control medium.

### Expression levels of select cell cycle and ECM genes are unchanged in ACF media

To explore any mRNA-level differences across media groups, RT-qPCR was used to examine a panel of commonly expressed genes important to cell cycle regulation, apoptosis, and extracellular matrix (ECM) and adhesion on NIH 3T3 fibroblasts grown in ACF media with different coatings, as compared to growth in FBS-containing medium. In total, 84 genes were assessed in a custom-made plate using Qiagen’s RT2 Profiler PCR Arrays. The full panel, their abbreviations, and fold changes with P-values can be found in the *Appendix, Table 1*. The selection of genes for the RT² Profiler PCR Array was driven by the overarching goal of understanding the fundamental molecular mechanisms underlying fibroblast function and cellular health. By integrating genes from Qiagen’s pre-made panels on ECM and adhesion, apoptosis, and cell cycle regulation, our custom panel aimed to comprehensively assess the molecular landscape governing fibroblast behavior. Fibroblasts orchestrate tissue integrity and function through their dynamic interplay with the ECM, their capacity to modulate cell-cell and cell-matrix interactions, and their pivotal roles in proliferation, survival, and response to environmental cues [89,90]. Thus, by interrogating genes associated with ECM/adhesion, apoptosis, and cell cycle regulation, our analysis aimed to provide comprehensive insights into the molecular underpinnings of normal fibroblast function, with implications for tissue homeostasis, repair processes, and potential therapeutic interventions.

After 30 days in culture, mRNA was harvested from all ACF media groups and the FBS group, where all steps necessary for RT-qPCR were carried out. During analysis, ACF media groups were compared to the FBS group, where genes with a >2-fold difference, a P-value of <0.05, and a C_T_ value cut off of 35 were deemed significantly changed from the FBS group, as represented in the clustergram (Fig. 6A). Volcano plots generated from this data exhibit these results for all genes studied (Fig. 6B-D). Of all genes in the panel, 54 genes for non-coated ACF group, 40 genes for ACF+gelatin group, and 39 genes for the ACF+soy group were found to be insignificantly different than the FBS group, represented by the black points in each volcano plot. To better understand some of the differences among the ACF media groups and why they may have caused NIH 3T3s to behave slightly differently from one another, we examined the significantly changed genes by categorizing them into functional groups - apoptosis (Fig. 6E), cell cycle (Fig. 6F), and ECM/adhesion (Fig. 6G). 7 genes relating to apoptosis were differentially expressed across all groups. Among these, 3 genes (BNIP3L, XIAP, AKT1) were downregulated (between 2- and 4-fold) across all groups. DAPK1 was up-regulated across all groups (between 5- and 6-fold) and IL10 was heavily up-regulated in gelatin (28.21-fold) and soy (12.12-fold) groups, with no change in expression in the non-coated group. 11 genes relating to cell cycle (CCNA2, SMC1A, CDC6, ATR, RBL1, BIRC5, CDKN3, CDC7, WEE1, AURKA, CHEK1) were down-regulated to a similar extent across all groups (between 2- and 5-fold). 8 genes relating to ECM/adhesion were differentially expressed. Up regulation of LAMA2 (ECM component) was present in all groups, although it showed highest expression in the non-coated group (13.86-fold) as compared to gelatin (2.26-fold) and soy (7.00-fold) groups. Consistent down-regulation of ITGA5 (integrin sub-unit) and CCN2 (matricellular protein) was present in all groups (between 3- and 4-fold). Other down-regulated genes showed varied expression across non-coated, gelatin, and soy groups: FN1 (−2.2-fold, −8.18-fold, −3.24-fold, respectively), SELP (−9.96-fold, −4.09-fold, −4.86-fold, respectively), FBLN1(−14.32-fold, −2.45-fold, −7.69-fold, respectively), and TGFBI (−4.5-fold, - 14.51-fold, −7.47-fold, respectively). Finally, among all differentially expressed genes relating to ECM/adhesion, only COL5A1 showed down-regulation among gelatin (−5.62-fold, P=0.0005) and soy (−3.61-fold, P=0.0007) groups, with non-coated groups remaining unchanged.

**Figure 6:**
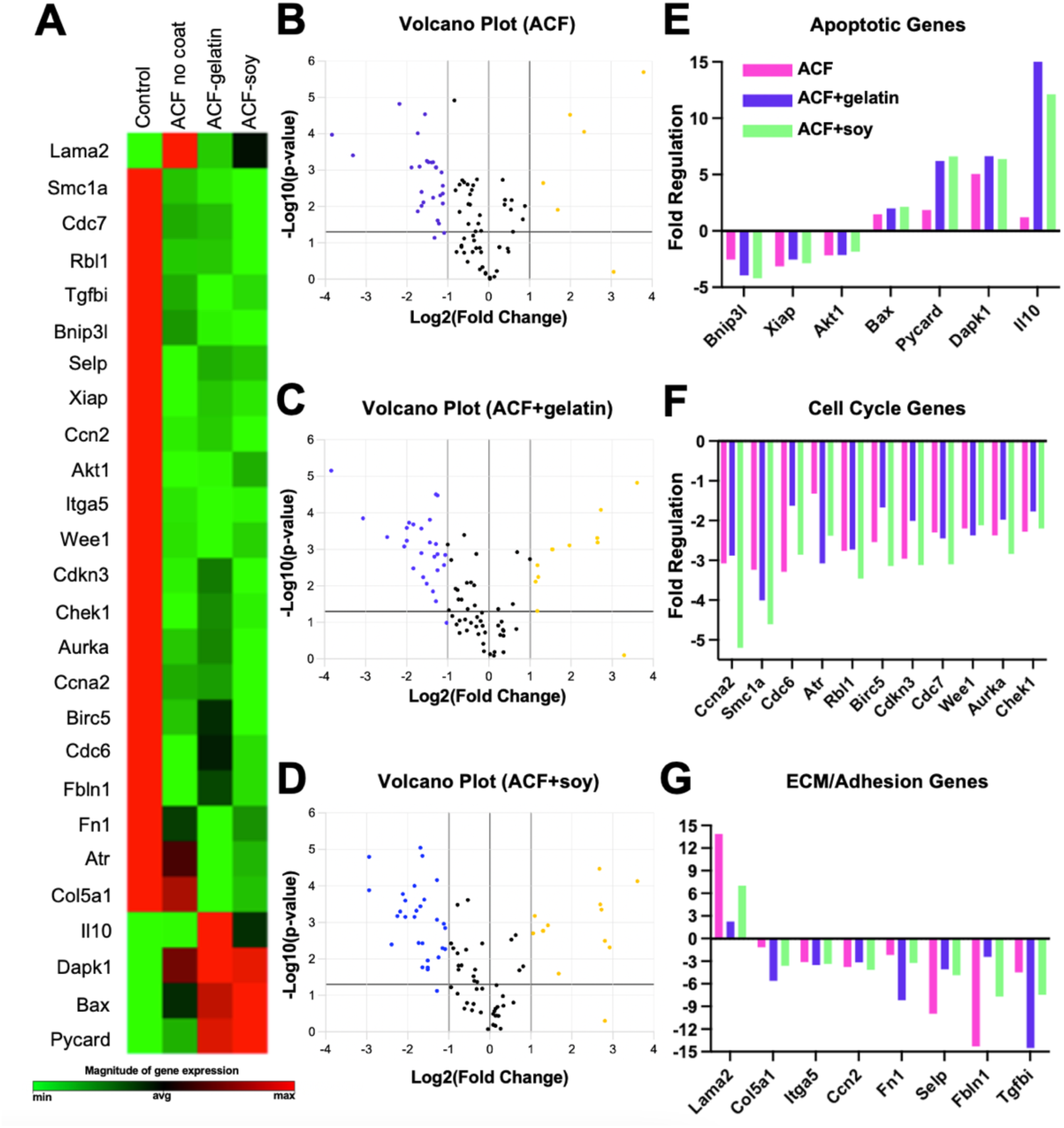
Differentially expressed genes between each media condition and the control through RT^2^ profiler PCR array analysis. (**A)** Clustergram of significantly altered genes (compared to the control group) from custom-designed gene panel across all groups. Each condition was analyzed in triplicate. Red represents maximum expression, green represents minimal expression, and black represents average levels of expression. **(B-D)** Volcano plots for each media condition. In these scatter plots, yellow points on the right side of the vertical line represent up-regulated gene expression as compared to the control group. Blue points on the left of the vertical line represent down-regulated gene expression as compared to the control group. Black points within the vertical lines represent unchanged gene expression, where fold change <2. Any points above the horizontal line at y=1.3 represent differentially expressed genes with a significant P-value <0.05. **(E-G)** Fold regulation of significantly differentially expressed genes across media conditions, grouped by function. The significant threshold for these genes was set as log2(fold change) > 2. This is compared with the control (FBS-based) group, for which fold regulation for each gene is normalized to 1. Significance values were defined at P<0.05.

### ACF media can support the growth of several adherent model cell lines

Although the ACF medium in this study was used to test its proliferative capacity on NIH 3T3 fibroblasts, we also selected other adherent cell lines to test the medium’s universal growth capabilities. In this set of experiments, C2C12 mouse myoblasts, HFF human fibroblasts, and MC3T3 mouse pre-osteoblasts were grown in both ACF+soy and FBS mediums and cultured for 30 days. It is important to note that the media formulations used for different cell types were slightly altered to meet each cell line’s base needs, reflected in Table 2. HFF fibroblasts were cultured in the medium identical to that used for NIH 3T3s, but for C2C12 and MC3T3 cell lines, a minor change in the basal medium, concentration of FGF2, and addition of TGFb was needed for optimized serum-free growth. On the 30th day of culture, mean cell concentrations were computed where each group showed a significantly increased total cell concentration in the ACF+soy group as compared to the FBS group (Fig. 7A). All cell types in ACF+soy medium were able to viably proliferate (C2C12s and MC3T3s: viability >94%, HFFs: viability >86%) as determined by Annexin-V/PI staining, with no significant changes in viability groups (Fig. 7B). In order to survey the confluency and general morphology of the cells at these time points, phase contrast images were obtained (Fig. 7C-H). All cell types grown in ACF+soy showed similar densities (>75%) to the FBS-based group. It is evident that the morphology (distinct clustering, cell elongation, specific characteristics) of each cell type was conserved when qualitatively comparing each image to its corresponding FBS-based control.

**Figure 7:**
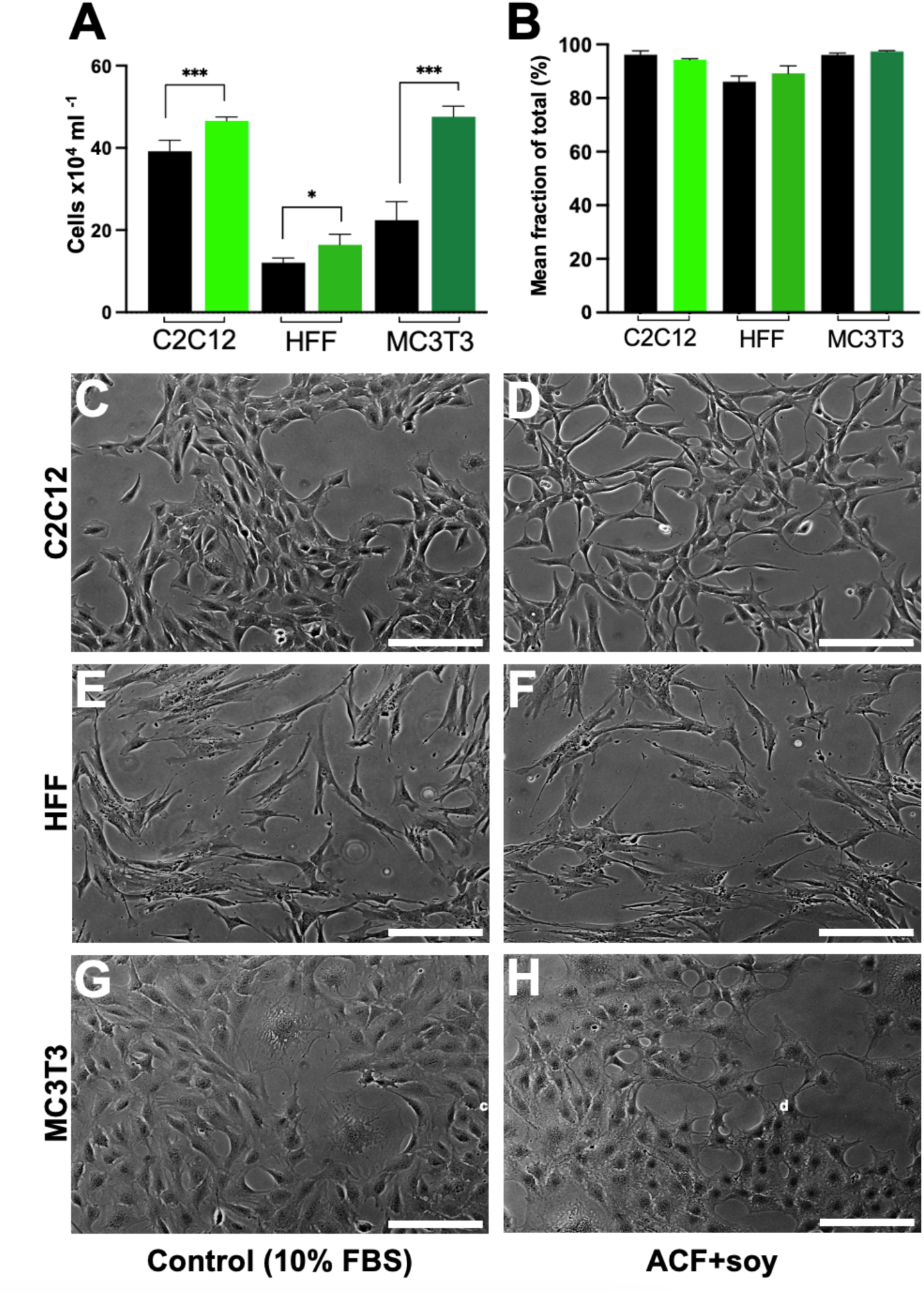
Cell concentration, viability, and morphology of 3 cell lines after 30 days. **(A)** Mean concentration of total cells (x10^4^ per ml). **(B)** Mean percentage of live cells as measured by Annexin-V/PI staining. N=3 for each cell line and each media condition. Each biological replicate is the mean of 3 technical replicates. Data are presented as mean ± SEM where *P<0.033, **P<0.002, ***P<0.001, as compared to the control (FBS-based) medium. **(C-H)** Most representative phase contrast images. Scale bars, 200um. Each row presents images for one cell line. **(C,E,G)** Control medium (10% FBS). **(D,F,H)** ACF medium (soy-coated).

## Discussion

Alternatives to fetal bovine serum have been continually emerging over the last few decades, reflecting a sustained effort to develop more controlled and defined culture conditions [91–93]. Previously fabricated SFM supplements serve as a robust foundation for creating a more universally applicable serum substitute but despite these commendable advancements, the growing reliance on FBS in various sectors, including industry, pharmaceutical manufacturing, cell culture, and research, underscores the need for a precisely regulated, well-defined serum-free alternative capable of accommodating diverse cell lines [94,95]. The research conducted in this study involves the thorough evaluation of an ACF culture system’s potential as a replacement for FBS. It encompasses an examination of overall cell viability and proliferation capacity for 4 commonly used adherent cell lines in research, as well as morphological and transcriptomic analyses for one model cell line, NIH 3T3 fibroblasts. A major point of this work is to not only validate a new ACF medium via a variety of techniques, but also to provide the cell culture community with a formulation that can be used on more than one cell line.

In our study, various concentrations of human or mouse recombinant growth factors and proteins that are typical of SFM development were combined and initially tested on fibroblasts. By conducting short-term (<7 day) qualitative and quantitative analyses, we were able to identify an ideal cocktail of components to be used in further experiments. We defined the ‘optimal’ formulation as one that was able to preserve cell viability and morphology for at least 7 days compared to the FBS condition and which contained a minimal number of components at the lowest effective concentrations. This optimization process was accomplished through a matrix which allowed us to test different concentrations of synergistic components (Table 1). Interestingly, our final formulation did not include transforming growth factor beta (TGFβ) and prostaglandin E2 (PGE2), which other SFM studies have often recommended and/or implemented themselves [23,96–99]. TGFβ is typically involved in regulating cell growth, proliferation, and differentiation [100,101], while PGE2 is known for its role in inflammation and cell signalling [102]. Our data suggested that their inclusion altered cell morphology and did not significantly enhance cell viability under our specific conditions. This indicates that the requirements for maintaining NIH 3T3 fibroblasts may differ from other cell types or conditions in other SFM studies. Although PGE2 was excluded for all cell lines, TGFb was still included in the growth of C2C12 myoblast and MC3T3 pre-osteoblast lines. Moreover, concentration dependencies played a crucial role in our findings. For example, epidermal growth factor (EGF) exhibited a dose-dependent effect where higher concentrations led to a more cobble-stone morphology as compared to the control. The minimal effective concentrations of components were critical not only for cost-effectiveness but also for mimicking typical morphologies of FBS-grown cells.

Cell adhesion in serum-free cultures has always been a factor of importance since FBS is the source of cell-adhesion promoting factors (e.g. vitronectin, fibronectin) to the culture system [23,92,103]. Thus, its absence often poses a problem for SFM development in adherent cell lines [32,104]. Although the adaptation of cells to suspension cultures or pre-coating substrate vessels with factors such as vitronectin or gelatin have been commonly employed solutions to such a problem, these solutions can be undesirable due to excess costs and their animal-derived nature. Significant research has been conducted employing media that incorporate plant-derived hydrolysates, as evidenced in prior studies [44,105–113]. These plant-based hydrolysates are often preferred over animal tissue-derived counterparts due to safety, ethical, and cost considerations. Soy protein hydrolysates have demonstrated a substantial capacity to enhance cell growth and the production of recombinant proteins in cell cultures and have been integrated into SFMs previously [106–113]. When using gelatin or soy protein hydrolysate as a substrate coating and then culturing the cell lines of focus in this study, the ACF medium showed enhanced growth rates in both short-term (<7 day) and long-term (90+ day) conditions. By assessing nuclear and cell body areas, we sought to capture the variations in morphology that may reflect underlying cellular responses to the altered culture environment. When fibroblasts were grown in the ACF medium without any substrate coating, nuclear to cell body ratios were significantly smaller than the FBS group. Without changing any components of the ACF medium but instead coating the cell culture substrates with either gelatin or soy protein hydrolysate solutions, nuclear to cell body ratios were rescued and found to be insignificantly different as compared to the FBS-grown cells. These findings suggest that soy protein hydrolysate is a crucial component to maintaining the adhesion and therefore the morphology of cells in this culture system, and that it may be a suitable plant-based substitute to gelatin coating in certain cell culture applications. Unlike previous studies that utilized combinations of FBS with soy protein hydrolysates, our approach eliminates all animal-derived components from the culture system, addressing both ethical concerns and cost-effectiveness more comprehensively. This complete removal of animal components reduces the risk of variability and contamination associated with animal-derived materials. Furthermore, by comparing the nuclear to cell body ratios under different substrate coatings, our findings highlight the essential role of soy protein hydrolysate in maintaining cell morphology and function. This contrasts with previous methodologies that often relied on animal-derived coatings, thus underscoring the novelty and practicality of our approach in creating a completely animal-free culture system.

In addition to proliferative and morphological forms of validation, we assessed the transcriptomic effects of this ACF medium on NIH 3T3 fibroblasts. The majority of genes relating to cell cycle, apoptosis, and ECM/adhesion remained unchanged, yet differentially expressed genes in the three serum-free groups, each with different coatings (non-coated, gelatin, and soy) revealed intricate molecular adaptations governing cell behavior. The complex processes of cell adhesion have a direct impact on the interactions between cells and their surrounding ECM, thereby influencing critical events like cell proliferation, differentiation, and tissue formation [114,115]. A shared response across all groups, regardless of coating type, was the upregulation of Laminin alpha 2 (LAMA2), a fundamental component of the basement membrane, crucial for cell adhesion, differentiation, and migration [116]. This upregulation present in all ACF groups indicates a concerted effort to fortify adhesion in serum-free conditions. Notably, the non-coated group exhibited the highest upregulation of Lama2 (13.86-fold compared to control), suggesting a compensatory effort to establish robust cell adhesion in the absence of FBS or any substrate coating. Gelatin is known to enhance cell adhesion via RGD motifs, a repeat amino acid sequence that is recognized by integrin receptors on cell surfaces, which when bound promote adhesion and other signalling cascades [53,54]. We speculate that RGD motifs present in Lunascin, a protein found within soy, may be replicating gelatin’s adhesive effects in culture which could also explain its compensatory effect in rescuing nuclear to cell body area ratios. However, it is also plausible that the soy protein hydrolysate may simply have altered the surface charge of the substrate, thereby promoting better adhesion through electrostatic interactions. Studies have shown that surface charge can significantly influence cell adhesion by affecting protein adsorption and integrin binding [117–119]. This change in surface charge could enhance cell attachment and spreading without necessarily involving specific adhesive motifs. The downregulation of Collagen Type V Alpha 1 (Col5a1) in the gelatin (−5.62) and soy-coated groups (−3.61) supports this notion. Collagen V is integral to the ECM, contributing to collagen fibril assembly [120]. The reduced expression of Col5a1 may indicate that cells in these coated conditions did not need to synthesize as much collagen, likely due to the external adhesive cues provided by the coatings, which facilitated cell attachment and maintained cell morphology.

Furthermore, the downregulation of fibronectin (FN1), transforming growth factor beta induced (TGFBI), cellular communication network factor 2 (CCN2), fibulin-1 (FBLN1), and integrin alpha 5 (ITGA5) in all groups signifies a collective deviation in cell-ECM interactions. This group of genes are all crucial components of the ECM that facilitate cell adhesion by interacting with integrins and other matrix proteins, promoting stability within the tissue microenvironment [121–124]. This downregulation, particularly pronounced in the non-coated group, implies a shift in cell-ECM interactions, potentially influencing cell attachment and migration capabilities. These findings correlate to the observation that cells in SFM tend to be less adherent, likely due to the lack of attachment factors that are typically present in FBS, typically like fibronectin [23,125,126].

Expanding our investigation to genes related to the cell cycle, a consistent downregulation of Cdc7, Cdkn3, Chek1, and Wee1 across all groups implies a collective suppression of cell cycle regulatory mechanisms in serum-free conditions. Cdc7 and Chek1 are crucial for DNA replication and repair, while Cdkn3 and Wee1 are involved in checkpoint control [127–130]. This downregulation likely reflects an adaptive response to the serum-free environment, where a slower cell cycle may help conserve resources and ensure cell survival under nutrient-limited conditions. The upregulation of Death-Associated Protein Kinase 1 (DAPK1) in all ACF groups indicates an increased sensitivity to apoptotic signals. DAPK1 is a pro-apoptotic kinase involved in mediating cell death in response to various stress signals [131]. This heightened apoptotic potential may have served as a protective mechanism, ensuring that damaged cells were efficiently removed, thus maintaining the overall health of the culture. Finally, the upregulation of PYCARD and Interleukin-10 (IL10) specifically in the gelatin and soy-coated groups reveals an interesting modulation of inflammatory responses. PYCARD is associated with inflammasome activation, leading to the production of pro-inflammatory cytokines [132], while IL10 is an anti-inflammatory cytokine that helps regulate immune responses and prevent excessive inflammation [133]. While cells grown in vitro in FBS are considered the standard for cell culture, they do not replicate the native in vivo environment. The observed changes in the balance of inflammatory and immune responses in serum-free conditions are not necessarily negative; rather, they reflect adaptations to a different culture environment. These adaptations result in similar outcomes in terms of morphology and cell viability, suggesting that neither condition is inherently better, but each supports cell health through different mechanisms.

While this study provides a promising resource for researchers and serves as a valuable foundation for ongoing media development, it’s evident that further optimization of media, focusing on cost-effectiveness and application to more cell lines, is necessary. The current approach in this work involved a step-by-step exploration of individual media components to identify suitable combinations for adherent cell lines. The interactions between these components can be complex and highly interdependent as they work in synergy to provide the necessary conditions for cells to grow and function properly. An in-depth understanding of the interplay between culture media components can be found in Yao and Asayama’s 2017 review [134]. Given the intricate interaction of nutrients, growth factors, hormones, buffering agents, and trace elements within cell cultures, it’s improbable that our soy-based ACF medium is already in its optimal form. There’s a possibility that certain modifications of our medium (e.g. without coating, with gelatin coating, or higher concentrations of soy coating) might provide advantages in untested cell lines or in 3D culture modalities (e.g. suspension cultures). Computational methods for media development are better equipped to address multifactorial challenges like this and should be utilized to advance serum-free media for various applications [135,136].

Several recent studies have proposed animal component-free media formulations, including the works of Mohr et al. (2023) and Schnell et al. (2024), which were developed for specific cell therapy and regenerative medicine applications [137,138]. These formulations share certain core components with ours - such as recombinant albumin, ITS, and defined growth factors, but they differ in their base media and coating strategies. For example, Schnell et al. employed a defined protein-free base with selective supplementation for mesenchymal stem cells, while our formulation uses a DMEM/F12 base enriched with recombinant human insulin, transferrin, albumin, ethanolamine, and variable supplementation with EGF, FGF-2, and TGF-β. Furthermore, our use of plant-derived soy hydrolysate or gelatin as a surface coating offers a modular approach to modulating cell adhesion in a manner not explored in those formulations. These differences underscore the diversity of strategies within the field and highlight the need for tailored optimization depending on the intended application and cell type. One limitation of the present study is the exclusive use of murine cell lines, which, while valuable for early-stage validation and benchmarking, may not fully reflect the clinical or translational potential of the ACF medium. To build on this work, future studies should investigate the medium’s performance in a broader range of human-derived cells, including primary and stem cell-based models. Moreover, while cells in this study were transitioned to ACF media following initial expansion in FBS-containing conditions, future research would benefit from direct isolation and long-term culture entirely under animal-free conditions, particularly for applications in regenerative medicine and biomanufacturing. Lastly, although animal-derived trypsin was used for passaging here to maintain consistency with standard protocols, we recommend the use of recombinant, animal-free alternatives such as TrypLE or TrypZean in subsequent studies to ensure a fully animal component-free workflow.

The research presented in this study introduces a completely animal component-free, soy-based medium to serve as a suitable replacement to FBS for NIH 3T3 cell culture in addition to other select cell lines (MC3T3, C2C12, HFF). However, achieving the universal nature of mammalian cell culture maintenance that FBS possesses, with the goal of preserving simplicity in the formulation, resemblance to typical cellular morphologies and behaviours, and cost-effectiveness, demands further refinement. Future endeavors to undertake these challenges might involve computational approaches using multifactorial analysis, machine learning algorithms, and other high throughput methods to better understand signalling pathways and the synergistic effects of different components in cell culture medium. Regardless, our results not only offer definitive support for using soy protein hydrolysate as a supplement to serum-free culture but also contribute to transparent methods in serum-free media development. The cellular proliferation rates in the soy-based medium proved to be at least as favorable, if not better than that in the conventional FBS-based medium. Morphological attributes of cells grown in the soy-based medium were matched to the FBS group and the majority of 84 genes studied by RT-qPCR remained unchanged. These findings have significant implications for refining cell culture methodologies, enhancing reproducibility, and tailoring culture conditions for diverse experimental settings.

## Methods

### Media formulation

We conducted a multi-step approach to explore different combinations of additives for cell culture media. We analyzed how these components interacted and adjusted their concentrations to develop a SFM recipe capable of supporting NIH 3T3 fibroblast growth. The optimization process is discussed in full in the Results. The final formulation with catalogue numbers and production sources can be found in Table 2 for brevity. A summary is included here, where these components were added to stock basal media in the following order:

1. Albumin (human recombinant): stock solution of 40mg/mL was created. A final concentration of 1mg/mL was used.
2. Insulin-transferrin-selenium (ITS, insulin/transferrin being human recombinant): Purchased as one supplement of stock solution 100X. A final solution of 1X was used, with final concentrations of 50 ug/mL (insulin), 27.5 ug/mL (transferrin), 25 ng/mL (selenium).
3. Ethanolamine: stock solution of 1013.93 mg/mL (16.6M) was created. A final concentration of 1.5mg/mL was used.
4. Fibroblast growth factor 2 (FGF-2, mouse recombinant): A stock solution of 100 μg/mL was created. A final concentration of 1 ng/mL (NIH 3T3, HFF cell types) or 30 ng/mL (C2C12, MC3T3 cell types) was used.
5. Epidermal growth factor (EGF, mouse recombinant): A stock solution of 10 ng/mL was created. A final concentration of 10 ng/mL was used.
6. Transforming growth factor beta (TGF-β, human recombinant): A stock solution of 100 μg/ml was created. A final concentration of 5 ng/mL was used (C2C12, MC3T3 cell types only).

### Cell lines and culture conditions

Murine embryonic fibroblast NIH 3T3 cells (ATCC-CRL-1658), murine MC 3T3 E1 Subclone 4 pre-osteoblast cells (ATCC CRL-2593), murine C2C12 myoblast cells (ATCC CRL-1772), and human foreskin fibroblast HFF cells (ATCC-SCRC-1041) were purchased from ATCC. For routine cell maintenance, cells were cultured at 37°C in 5% CO2, passaged using 0.05% trypsin-EDTA (Cytiva, SH3023602), and counted using trypan blue with an image-based cell counter (Countess^TM^ II FL, Invitrogen). Initially, NIH 3T3s and HFFs were maintained in proliferation medium consisting of 1:1 DMEM/Ham’s F-12 (Sigma, 51445C) supplemented with 10% (v/v) FBS (Wisent, cat. 098150, lot. 185729) and 1% (v/v) penicillin/streptomycin (P/S; 100 U/mL penicillin, 100 mg/ml streptomycin; Cytiva, SV30010). The same protocol was used for the maintenance of MC3T3 cells and C2C12 cells but with different basal mediums - MEM Alpha 1X (Gibco, A1049001) and DMEM (Sigma, D0822), respectively.

To maintain consistency, the medium used to culture the FBS-grown groups contains the same basal medium but is supplemented with 10% FBS (Wisent, cat. 098150, lot. 185729). It is also important to note that this one lot of FBS has been secured for the entire duration of this study, to minimize variability across FBS-grown samples.

Cells were maintained in 10 cm culture dishes at a seeding density of 8.8×10^3^ cells/cm^2^ and sub-cultured every 2 days at around 85% confluence. After 2 passages in this serum-containing medium, cells were either kept in this same medium, designated to be controls grown in serum-containing medium, or went on to undergo the serum adaptation process, designated to eventually become the treatment groups grown in 100% serum-free medium.

### Adaptation and serum-free media

Serum adaptation was completed through five sequential passages over ten days, to diminish any stress the serum switch may have caused on the cells of the treatment groups. This adaptation process was repeated every round of media optimization before other validation. The cells in the FBS groups did not undergo the serum adaptation process, although they were sub-cultured every 48 hours in serum-based medium during this time in order to mature to the same passage number as the treatment groups. All FBS groups and all treatment (ACF) groups were individually cultured in triplicate in 10 cm culture dishes at seeding densities of 8.8×10^3^ cells/cm^2^. All sub-culture procedures are identical to those explained above, other than the ACF medium used in the adaptation process for the treatment groups. Cells of treatment groups were initially cultured in 100% FBS-based medium and after 48 hours were then sub-cultured in medium consisting of 80% FBS-based medium and 20% ACF medium. Every 48 hours, as the cells reached around 85% confluency, they were sub-cultured to sequentially reduced serum amounts, with 60% FBS-based medium and 40% ACF medium, followed by 40% FBS-based medium and 60% ACF medium, 20% FBS-based medium and 80% ACF medium, and finally 100% ACF medium. In the following sections, day 0 for each treated group is defined as 1 passage post seeding cells in 100% ACF medium.

Treatment groups of NIH 3T3s were adapted and routinely cultured in a variety of in-house developed ACF culture media formulations. These formulations are composed of DMEM/Ham’s F-12 (Sigma, 51445C) with 1% (v/v) P/S as its base, to match the FBS group’s medium base. They are supplemented with a variety of hormones, carrier proteins, lipids, minerals, and growth factors. The final iteration of the ACF medium’s components and its corresponding concentrations can be found in Table 2. Basal media is the foundational liquid medium that provides essential nutrients, pH buffers, and other necessary components to support the growth of cells in vitro [91]. It is often supplemented with antibiotics, animal sera, and/or growth factors to create a specific environment tailored to the needs of the cells being cultured. The reason that DMEM/F-12 medium was chosen as the basal medium is because when traditional mediums (i.e. DMEM, MEM, Ham’s F-12) were used in serum-free cultures, their ability to maintain growth was often inadequate, yet a 1:1 ratio of DMEM and Ham’s F-12 exhibited better performance when used for serum-free culture of certain cell types [139,140]. As a result, DMEM/F-12 basal media has been recommended by a variety of studies aiming to create serum-free media formulations [14,18,19]. In this study, this basal medium combined with ACF supplements was ideal for growing NIH 3T3 fibroblasts and HFF human fibroblasts, but C2C12 myoblasts and MC3T3 pre-osteoblasts were grown in other basal mediums that are typically recommended for their growth in standard protocols (DMEM and aMEM, respectively). In the FBS groups for these cell lines, the same basal medium is used as in the ACF-supplemented groups.

### Animal- and plant-based substrate coatings

SFM alternatives often incorporate supplementary additives to facilitate cell adhesion, compensating for the absence of components typically provided by FBS, thereby ensuring optimal cell growth and viability [52,141–143]. Such additives are often animal-derived and include extracellular matrix components such as fibronectins, vitronectins, laminins, and collagen [52,143–145]. A notable approach in achieving animal-free conditions has involved incorporating plant-based components as viable alternatives [91,146]. These components commonly include extracts from wheat, soy, rice, chickpea, pea, and other protein-rich botanical sources and have shown to enhance cell proliferation and improve recombinant protein production in some mammalian cell lines [44,105,106,147]. Initially, a gelatin coating on the substrate was used to increase cell adhesion and to observe if it would have an effect on the poor attachment seen in the preliminary SFM we created. Through researching the literature, soy protein hydrolysate was identified as a substitute to gelatin coating in this context.

Renowned for its rich nutritional profile and diverse bioactive compounds, soybean stands out as a promising candidate to promote cell viability, proliferation, and aiding in adhesion [107,148]. Soy protein hydrolysates generated through diverse methods have outperformed other plant-based hydrolysates in enhancing cell growth in a diverse set of cell lines and culture applications [44,106–113]. Via further validation, soy as a coating was deemed an essential piece to the success of our final ACF media. Gelatin coatings were produced by dissolving gelatin (Sigma, G9136) in DI H2O at a concentration of 1 g/L. Soy coatings were produced by dissolving soy protein hydrolysate (Sigma, S1674) in DI H2O at a concentration of 4 g/L. Gelatin and soy solutions were autoclaved once to sterilize. 24 hours before cell seeding, 10 cm dishes were each coated with 8 mL of soy or gelatin solutions. Right before cell seeding, solutions were aspirated from each dish and corresponding media was carefully added to not disrupt the particles adsorbed to the surface.

### Proliferation rates and viability

The passage number for all groups and their replicates in these studies was P8 at day 0. Day 0 of short- and long-term counts is designated as 1 passage after the treatment groups were placed in 100% ACF medium, post-10-day serum adaptation process. At this point, FBS and ACF groups were sub-cultured with their corresponding media and inoculated with seeding densities pertaining to either short-, or long-term counts, described in detail in the following sections. For each timepoint and each group (FBS or ACF), 3 biological replicates were analyzed. 3 technical replicate values were obtained for each biological replicate in every condition.

All FBS and ACF groups were inoculated with a seeding density of 1×10^4^ cells/mL per culture plate on day 0. All conditions were monitored for this short-term growth period, with readings taken after 1, 4, and 7 days of culture. At each time point, cells were trypsinized and centrifuged at 1000 rpm (97g) for 3 minutes. After supernatant removal, cell pellets were re-suspended in 2 mL of PBS before being treated with the Annexin V-FITC Apoptosis/Staining Detection Kit (Abcam, ab14085). Briefly, for each sample, 1000 mL of 1 x binding buffer was added to the cell pellet after centrifugation and PBS removal. After re-suspension in binding buffer, Annexin V-FITC (5 mL) and PI (5 mL) were added to 100 mL of cell suspension. After 15 minutes of incubation at room temperature in the dark, 400 mL of 1 x binding buffer was added, resulting in a final volume of 500 mL for each sample. Keeping samples on ice in the dark, all samples were counted using the Countess^TM^ II FL cell counter equipped with fluorescence filters (Ex 482/25, Em 524/24; EVOS Light Cube for GFP; Invitrogen, AMEP4951) and (Ex 531/40, Em 593/40; EVOS Light Cube for RFP; Invitrogen, AMEP4952). To avoid false recognition of debris or evident noise by the cell counter, an arbitrary size threshold level was set (size of 2–30 um) along with the corresponding fluorescence intensity (relative intensity of 40–255) for each assay. Each threshold level was confirmed by reference to past studies of NIH 3T3 cell diameter and by examination of the real-time detection image of each cell using the counter.

For long-term growth studies (90-day – NIH 3T3; or 30-day – other cell types), all FBS and ACF groups were inoculated with a seeding density of 5×10^4^ cells/mL per culture plate on day 0. All groups were monitored for a period of 90+ days or 30 days in triplicate. During this period, subcultures for each dish were performed every 72 hours, where cell viability was quantified using trypan blue and the automated cell counter. Live cell counts were recorded for each replicate and then inoculated at a concentration of 5×10^4^ cells/mL in new dishes at each subculture. Cell viability was quantified at 30, 60, and 90 days of culture (shown in this study), and 120, 150, 180, and 210 days of culture (data not shown) via the Annexin V-FITC Apoptosis/Staining Detection Kit in the same manner as described above.

### Fluorescent staining and microscopy

At day 0, cells for each condition were seeded in 35 mm dishes at a concentration of 1×10^4^ cells/mL. After 7 days, cells were washed three times with 1 x PBS. The samples were fixed in 4% paraformaldahyde (PFA) for 10 min and then washed with PBS three times. Following the final PBS wash, cell permeabilization was achieved via addition of room temperature 0.02% TritonX-100 for 3 min. F-actin was stained using 1:200 Alexa Fluor^TM^ 546 Phalloidin (Invitrogen, A22283). Nuclei were visualized using 1:500 DAPI (Invitrogen, D1306). Phalloidin and DAPI were added together in PBS at their respective concentrations and then incubated for 1 hour in the dark at room temperature. Following 3 washes with PBS, Vectashield mounting medium was added to each sample. A round glass coverslip was carefully placed on the surface of the dish, and samples were visualized by confocal laser scanning microscopy through the coverslip by inversion of the 35 mm dish on an AFM insert. Confocal imaging was performed on an A1R high speed laser scanning confocal system on a TiE inverted optical microscope platform (Nikon, Canada) with appropriate laser lines and filter sets.

Image processing was executed via ImageJ open access software using a semi-automatic method. Briefly, DAPI (blue) and Phalloidin (red) channels were split. Next, Z-stacked 3D volume images were prepared for each channel. For the DAPI channels, a threshold was set, converting the colored max intensity projection to a binary image. Next, the Analyze Particles plugin was used to measure each nucleus’ area, perimeter, circularity, roundness, aspect ratio, and other measures. If nuclei in the binary image were slightly warped and not fully picked up by the thresholding algorithm, that nucleus was excluded from data analysis. For the Phalloidin channels, each cell body was manually selected using the polygon selection tool and then measured within the software. The same measurements obtained from the nuclei were acquired. DAPI channels and Phalloidin channels were set side by side when performing the manual analysis on the cell bodies to ensure the order in which the cell bodies were counted would match the corresponding nucleus. This allowed for proper computation of each nucleus area to cell body area (N:C) ratios. Cell bodies of the Phalloidin channels were excluded if they were found on the bounds of the image or if they were obscured by other cells to accurately compute their area. 30 images were analyzed for each media group with included cells as follows: FBS group (619 cells included in analysis), ACF no coat group (671 cells included in analysis), ACF gelatin group (477 cells included in analysis), and ACF soy group (493 cells included in analysis). Beeswarm plots were created in GraphPad Prism, version 10.2.3 for MacOsx (GraphPad Software, La Jolla, CA, USA).

Transmitted light images were acquired on an inverted TiE microscope (Nikon, Canada) with phase contrast optics. Images were analyzed using ImageJ open access software (http://rsbweb.nih.gov/ij/). Brightness and contrast adjustments were the only manipulations performed to images.

### Gene expression and analysis

For NIH 3T3s, 30 days after sequential serum adjustment, gene expression analysis was performed for each biological replicate in both FBS and ACF groups. Each plate of cells was trypsinized, centrifuged at 1000 rpm (97g) for 3 minutes, and re-suspended in 1 mL of appropriate medium. Next, the cell suspension was centrifuged, washed with PBS, and centrifuged once again. Supernatant was collected, leaving only the cell pellet. Purification of total RNA was performed using the RNeasy Mini Kit (Qiagen, 74104) for animal cells using spin technology. cDNA was reverse transcribed from 2 mg total RNA from each sample, using the Qiagen RT2 First Strand Kit (Qiagen, 330404) following the manufacturer’s protocols. qPCR was performed using a custom-made RT2 profiler PCR array (Qiagen, 330171) following the manufacturer’s protocols. A full list of the gene panel can be found in Table 1 of the Supplement. Reactions were performed on a Bio-Rad CFX96 Real-Time PCR Detection System (Bio-Rad Laboratories, USA) and fold changes in gene expression were calculated via the DDCT method. Normalization of the raw data was performed with species-specific housekeeping genes B2m and Gusb using the geometric mean as the normalization factor. During qPCR analysis, C_T_ cut-off value used was 35. The primers chosen in the customized plate included 32 primers for cell cycle genes, 25 for apoptotic genes, 27 for extracellular matrix and adhesion genes, and 5 for housekeeping genes. This gene expression panel was carried out in each of the following groups, each with 3 biological replicates: control (FBS) group, treatment group 1 (ACF medium without coating), treatment group 2 (ACF medium with gelatin coating), and treatment group 3 (ACF medium with soy coating).

### Statistical analysis

A two-way analysis of variance (ANOVA) was applied to assess the changes in cell viability across media groups and across time. Using post-hoc tests (Dunnet’s multiple comparisons test), the specific differences between media groups at each time point were explored. The mean of all the biological replicates for each treated group was compared to the mean of the replicates for the FBS group. A one-way ANOVA using Dunnet’s method for multiple comparisons was applied to analyse the nuclear to cell area ratios across media groups. Differences were considered statistically significant with *P<0.033, **P<0.002, ***P<0.001. Statistical analyses for RT-qPCR were provided by Qiagen’s online analysis software where P<0.05 was considered statistically significant. Captions for figures and tables describing experimental results state the number of biological replicates (N) as mean ± SEM, unless otherwise stated. The results were analyzed using GraphPad Prism, version 10.2.3 for MacOsx (GraphPad Software, La Jolla, CA, USA).

## Data Availability Statement

Data available on request.

