## Supplemental Table for "The creation and validation of a fully animal component-free media for select adherent cell types"

**Appendix Table 1. Gene panel used in RT2 PCR Profiler Array.** Entire panel was purchased in a kit from Qiagen. Fold-Change (2^(- Delta Delta CT)) is the normalized gene expression (2^(-Delta CT)) in the Test Sample (ACF conditions) divided the normalized gene expression (2^(-Delta CT)) in the Control Sample (FBS-grown cells). Special notes can be given either *A* or *B*. *A* signifies that the gene's average threshold cycle is relatively high (>30), so its relative expression level is low in both control and test samples, and the p-value for the fold change is either unavailable or quite high (p>0.05). *B* means that this gene's average threshold cycle is either not determined or greater than the defined cut-off (Ct>35), meaning that the expression was undetected and therefore results are un-interpretable.

| Gene Name Abbreviation | Gene Name | NCBI Gene ID | Fold change (ACF), p-value, special note | Fold change (ACF+gelatin), p-value, special note | Fold change (ACF+soy), p-value, special note |
|---|---|---|---|---|---|
| Abl1 | c-abl oncogene 1, non-receptor tyrosine kinase | 11350 | 0.47<br>P=0.0529 | 0.69<br>P=0.1665 | 0.58<br>P=0.0896 |
| Cdc6 | cell division cycle 6 | 23834 | 0.30<br>P=0.0136 | 0.61<br>P=0.0890 | 0.35<br>P=0.0194 |
| Cdkn2b | cyclin-dependent kinase inhibitor 2B | 12579 | 1.20<br>P=0.1310<br>B | 2.92<br>P=0.0010<br>B | 2.46<br>P=0.0017<br>B |
| Rad17 | RAD17 checkpoint clamp loader component | 19356 | 1.10<br>P=0.8512 | 1.59<br>P=0.1514 | 1.49<br>P=0.1583 |
| Aifm1 | apoptosis-inducing factor, mitochondrion-associated 1 | 26926 | 0.63<br>P=0.1190 | 0.54<br>P=0.0666 | 0.52<br>P=0.0595 |
| Casp7 | caspase 7 | 12369 | 0.95<br>P=0.7077 | 2.27<br>P=0.0027 | 1.76<br>P=0.0159 |
| Mcl1 | myeloid cell leukemia sequence 1 | 17210 | 0.46<br>P=0.0082 | 0.84<br>P=0.2096 | 0.59<br>P=0.0196 |
| Xiap | X-linked inhibitor of apoptosis | 11798 | 0.32<br>P=0.0095 | 0.39<br>P=0.0143 | 0.35<br>P=0.0111 |
| Ctnnb1 | catenin (cadherin associated protein), beta 1 | 12387 | 0.71<br>P=0.0302 | 0.7<br>P=0.0429 | 0.62<br>P=0.0150 |
| Itgb4 | integrin beta 4 | 192897 | 0.78<br>P=0.0859 | 1.5<br>P=0.0316 | 1.25<br>P=0.1048 |
| Thbs3 | thrombospondin 3 | 21827 | 0.38<br>P=0.0062 | 0.32<br>P=0.0003 | 0.33<br>P=0.0002 |

| | | | | | |
|---|---|---|---|---|---|
| Hsp90ab1 | Melanocortin 1 receptor | | 0.41<br>P=0.0008 | 0.53<br>P=0.0020 | 0.46<br>P=0.0011 |
| Atr | ataxia telangiectasia and Rad3 related | 245000 | 0.76<br>P=0.1776 | 0.33<br>P=0.0058 | 0.42<br>P=0.0011 |
| Cdc7 | cell division cycle 7 | 12545 | 0.43<br>P=0.0297 | 0.41<br>P=0.0263 | 0.32<br>P=0.0170 |
| Cdkn3 | cyclin-dependent kinase inhibitor 3 | 72391 | 0.34<br>P=0.00003 | 0.5<br>P=0.0007 | 0.32<br>0.00002 |
| Rb1 | retinoblastoma 1 | 19645 | 1.02<br>P=0.9724 | 0.74<br>P=0.1028 | 0.86<br>P=0.2926 |
| Akt1 | thymoma viral proto-oncogene 1 | 11651 | 0.46<br>P=0.0027 | 0.47<br>P=0.0027 | 0.54<br>P=0.0051 |
| Casp8 | caspase 8 | 12370 | 0.88<br>P=0.0470 | 0.9<br>P=0.1244 | 0.76<br>P=0.0298 |
| Naip1 | NLR family, apoptosis inhibitory protein 1 | 17940 | 1.2<br>P=0.1310<br>B | 2.92<br>P=0.0010<br>B | 2.46<br>P=0.0017<br>B |
| Adamts5 | a disintegrin-like and metallopeptidase (reprolysin type) with thrombospondin type 1 motif, 5 (aggrecanase-2) | 23794 | 1.46<br>P=0.0122 | 0.79<br>P=0.0342 | 1.55<br>P=0.0022 |
| Fbln1 | fibulin 1 | 14114 | 0.07<br>P=0.0001 | 0.41<br>P=0.0007 | 0.13<br>P=0.0001 |
| Lama1 | laminin, alpha 1 | 16772 | 1.20<br>P=0.1310<br>B | 2.92<br>P=0.0010<br>B | 3.2<br>P=0.0252 |
| Tgfbi | transforming growth factor, beta induced | 21810 | 0.22<br>P=0.00002 | 0.07<br>P=0.00001 | 0.13<br>P=0.00002 |
| MGDC | Mouse genomic DNA control | N/A | N/A | N/A | N/A |
| Aurka | aurora kinase A | 20878 | 0.42<br>P=0.0242 | 0.51<br>P=0.0464 | 0.35<br>P=0.0172 |
| Cdk1 | cyclin-dependent kinase 1 | 12534 | 8.34<br>P=0.6282 | 9.79<br>P=0.7914 | 7.00<br>P=0.04964 |
| Chek1 | checkpoint kinase 1 | 12649 | 0.44<br>P=0.0049 | 0.57<br>P=0.0118 | 0.45<br>P=0.0050 |

| | | | | | |
|---|---|---|---|---|---|
| Rbl1 | retinoblastoma-like 1 (p107) | 19650 | 0.36 P=0.0006 | 0.37 P=0.0006 | 0.29 P=0.0005 |
| Bad | BCL2-associated agonist of cell death | 12015 | 0.7 P=0.0025 | 0.66 P=0.0004 | 0.69 P=0.0002 |
| Dapk1 | death associated protein kinase 1 | 69635 | 5.06 P=0.0001 | 6.62 P=0.0001 | 6.37 P=0.00003 |
| Nfkb1 | nuclear factor of kappa light polypeptide gene enhancer in B cells 1, p105 | 18033 | 0.61 P=0.0025 | 0.74 P=0.0080 | 0.58 P=0.0003 |
| Ccn2 | connective tissue growth factor | 14219 | 0.27 P=0.0008 | 0.32 P=0.0013 | 0.24 P=0.0007 |
| Fn1 | fibronectin 1 | 14268 | 0.45 P=0.0011 | 0.12 P=0.0001 | 0.31 P=0.0004 |
| Lama2 | laminin, alpha 2 | 16773 | 13.86 P=0.000002 | 2.26 P=0.0490 | 7.00 P=0.0032 |
| Vcan | versican | 13003 | 0.73 P=0.0089 | 0.28 P=0.0002 | 0.67 P=0.0056 |
| RTC | Reverse Transcription Control | N/A | N/A | N/A | N/A |
| Bcl2 | B cell leukemia/lymphoma 2 | 12043 | 1.51 P=0.0492 | 1.10 P=0.6465 | 1.65 P=0.0205 |
| Cdk2 | cyclin-dependent kinase 2 | 12566 | 0.40 P=0.0724 | 0.49 P=0.1027 | 0.41 P=0.0750 |
| Ddit3 | DNA-damage inducible transcript 3 | 13198 | 1.38 P=0.1427 | 2.20 P=0.0076 | 2.08 P=0.0020 |
| Shc1 | src homology 2 domain-containing transforming protein C1 | 20416 | 0.71 P=0.0086 | 0.87 P=0.0911 | 0.80 P=0.0666 |
| Bax | BCL2-associated X protein | 12028 | 1.48 P=0.0067 | 2.00 P=0.0911 | 2.14 P=0.067 |
| Diablo | diablo, IAP-binding mitochondrial protein | 66593 | 0.64 P=0.0019 | 0.69 P=0.0082 | 0.78 P=0.0197 |

| | | | | | |
|---|---|---|---|---|---|
| Nol3 | nucleolar protein 3 (apoptosis repressor with CARD domain) | 78688 | 1.15<br>P=0.5931<br>A | 1.28<br>P=0.2333<br>A | 1.10<br>P=0.6835<br>A |
| Cdh1 | cadherin 1 | 12550 | 1.40<br>P=0.1760<br>A | 2.92<br>P=0.0010<br>B | 2.46<br>P=0.0017<br>B |
| Itga2 | integrin alpha 2 | 16398 | 1.2<br>P=0.1310<br>B | 3.90<br>P=0.0008 | 2.46<br>P=0.0017<br>B |
| Lamb2 | laminin, beta 2 | 16779 | 0.84<br>P=0.0295 | 0.91<br>P=0.3842 | 0.77<br>P=0.0398 |
| Vtn | vitronectin | 22370 | 0.86<br>P=0.5341<br>A | 1.09<br>P=0.8141<br>A | 0.97<br>P=0.8387<br>A |
| RTC | Reverse Transcription Control | N/A | N/A | N/A | N/A |
| Birc5 | baculoviral IAP repeat-containing 5 | 11799 | 0.39<br>P=0.0057 | 0.60<br>P=0.0234 | 0.32<br>P=0.0037 |
| Cdk6 | cyclin-dependent kinase 6 | 12571 | 0.95<br>P=0.6706 | 1.27<br>P=0.1614 | 1.16<br>P=0.3650 |
| Dst | dystonin | 13518 | 1.79<br>P=0.225 | 2.30<br>P=0.0057 | 2.67<br>P=0.0012 |
| Skp2 | S-phase kinase-associated protein 2 (p45) | 27401 | 0.57<br>P=0.0036 | 0.48<br>P=0.0014 | 0.47<br>P=0.0014 |
| Bcl2l1 | BCL2-like 1 | 12048 | 0.90<br>P=0.4653 | 1.28<br>P=0.1705 | 1.26<br>P=0.1930 |
| Fas | nan | nan | 2.52<br>P=0.0022 | 12.18<br>P=0.00002 | 7.59<br>P=0.0048 |
| Pycard | PYD and CARD domain containing | 66824 | 1.85<br>P=0.0096 | 6.21<br>P=0.0005 | 6.60<br>P=0.0004 |
| Col1a1 | collagen, type I, alpha 1 | 12842 | 1.31<br>P=0.0088 | 0.82<br>P=0.0709 | 1.10<br>P=0.2318 |
| Itga4 | integrin alpha 4 | 16401 | 1.20<br>P=0.1310<br>B | 2.92<br>P=0.0010<br>B | 2.46<br>P=0.0017<br>B |
| Lamc1 | laminin, gamma 1 | 226519 | 0.77<br>P=0.0635 | 0.40<br>P=0.0016 | 0.52<br>P=0.0038 |
| Actb | actin, beta | 11461 | 0.30<br>P=0.00001 | 0.37<br>P=0.0002 | 0.28<br>P=0.0001 |

| | | | | | |
|---|---|---|---|---|---|
| RTC | Reverse Transcription Control | N/A | N/A | N/A | N/A |
| Casp3 | caspase 3 | 12367 | 1.02 P=0.9093 | 1.24 P=0.0958 | 1.14 P=0.2070 |
| Cdkn1a | cyclin-dependent kinase inhibitor 1A (P21) | 12575 | 0.70 P=0.3321 | 1.28 P=0.6869 | 1.20 P=0.8115 |
| Itgb1 | integrin beta 1 (fibronectin receptor beta) | 16412 | 0.66 P=0.1863 | 0.58 P=0.1139 | 0.74 P=0.2562 |
| Smc1a | structural maintenance of chromosomes 1A | 24061 | 0.31 P=0.0008 | 0.25 P=0.0006 | 0.22 P=0.0005 |
| Bid | BH3 interacting domain death agonist | 12122 | 0.66 P=0.0759 | 0.67 P=0.0834 | 0.76 P=0.1716 |
| Igf1r | insulin-like growth factor I receptor | 16001 | 0.61 P=0.0140 | 0.63 P=0.0237 | 0.59 P=0.0153 |
| Ripk1 | receptor (TNFRSF)-interacting serine-threonine kinase 1 | 19766 | 0.80 P=0.0483 | 0.59 P=0.0129 | 0.58 P=0.0070 |
| Col4a1 | collagen, type IV, alpha 1 | 12826 | 1.32 P=0.0065 | 0.94 P=0.1874 | 1.06 P=0.3228 |
| Itga5 | integrin alpha 5 (fibronectin receptor alpha) | 16402 | 0.32 P=0.0040 | 0.28 P=0.0033 | 0.30 P=0.0036 |
| Selp | selectin, platelet | 20344 | 0.10 P=0.0004 | 0.24 P=0.0008 | 0.21 P=0.0007 |
| B2m | beta-2 microglobulin | 12010 | 1.27 P=0.0018 | 0.89 P=0.0544 | 1.12 P=0.0525 |
| PPC | Positive PCR Control | N/A | N/A | N/A | N/A |
| Ccna2 | cyclin A2 | 12428 | 0.33 P=0.0077 | 0.35 P=0087 | 0.19 P=0.0040 |
| Cdkn1b | cyclin-dependent kinase inhibitor 1B | 12576 | 0.58 P=0.1778 | 0.61 P=0.1952 | 0.65 P=0.2294 |
| Mcm2 | minichromosome maintenance complex component 2 | 17216 | 0.35 P=0.0006 | 0.26 P=0.0002 | 0.23 P=0.0002 |
| Trp53 | transformation related protein 53 | 22059 | 0.86 P=0.0173 | 1.02 P=0.7390 | 1.08 P=0.2704 |

| | | | | | |
|---|---|---|---|---|---|
| Birc2 | baculoviral IAP repeat-containing 2 | 11797 | 0.66<br>P=0.0021 | 1.61<br>P=0.0012 | 1.44<br>P=0.0030 |
| Il10 | interleukin 10 | 16153 | 1.20<br>P=0.1310<br>B | 28.21<br>P=0.0067 | 12.12<br>0.0001 |
| Traf2 | TNF receptor-associated factor 2 | 22030 | 0.88<br>P=0.1782 | 1.17<br>P=0.1209 | 1.10<br>P=0.3597 |
| Col5a1 | collagen, type V, alpha 1 | 12831 | 0.86<br>P=0.1383 | 0.18<br>P=0.0005 | 0.28<br>P=0.0007 |
| Itgav | integrin alpha V | 16410 | 0.81<br>P=0.1390 | 0.81<br>P=0.1297 | 1.12<br>P=0.3565 |
| Sgce | sarcoglycan, epsilon | 20392 | 3.23<br>P=0.0122 | 2.92<br>P=0.0010<br>B | 2.46<br>P=0.0017<br>B |
| Gapdh | glyceraldehyde-3-phosphate dehydrogenase | 14433 | 0.56<br>P=0.00001 | 0.41<br>P=0.00003 | 0.31<br>P=0.00001 |
| PPC | Positive PCR Control | N/A | N/A | N/A | N/A |
| Ccne1 | cyclin E1 | 12447 | 1.02<br>P=0.8925 | 1.21<br>P=0.2207 | 1.06<br>P=0.6413 |
| Cdkn2a | cyclin-dependent kinase inhibitor 2A | 12578 | 1.20<br>P=0.1310<br>B | 2.92<br>P=0.0010<br>B | 2.46<br>P=0.0017<br>B |
| Notch2 | nan | nan | 0.63<br>P=0.1289 | 0.92<br>P=0.6166 | 0.68<br>P=0.1613 |
| Wee1 | WEE 1 homolog 1 (S. pombe) | 22390 | 0.46<br>P=0.0044 | 0.42<br>P=0.0037 | 0.47<br>P=0.0054 |
| Bnip3l | BCL2/adenovirus E1B interacting protein 3-like | 12177 | 0.39<br>P=0.0006 | 0.25<br>P=0.0003 | 0.24<br>P=0.0003 |
| Mapk1 | mitogen-activated protein kinase 1 | 26413 | 0.73<br>P=0.0046 | 0.78<br>P=0.0096 | 0.81<br>P=0.0426 |
| Trp53bp2 | transformation related protein 53 binding protein 2 | 209456 | 0.77<br>P=0.0026 | 0.81<br>P=0.0013 | 0.88<br>P=0.0038 |
| Ctnna1 | catenin (cadherin associated protein), alpha 1 | 12385 | 0.41<br>P=0.0009 | 0.42<br>P=0.0014 | 0.41<br>P=0.0008 |
| Itgb3 | integrin beta 3 binding protein (beta3-endonexin) | 67733 | 3.98<br>P=0.00003 | 6.26<br>P=0.0006 | 6.47<br>P=0.0003 |

| | | | | | |
|---|---|---|---|---|---|
| Sparc | secreted acidic cysteine rich glycoprotein | 20692 | 0.82<br>P=0.0018 | 0.42<br>P=0.0003 | 0.41<br>P=0.00007 |
| Gusb | glucuronidase, beta | 110006 | 0.75<br>P=0.0035 | 1.17<br>P=0.0433 | 0.88<br>P=0.0656 |
| PPC | Positive PCR Control | N/A | N/A | N/A | N/A |
